# Lipid dependence of connexin-32 gap junction channel conformations

**DOI:** 10.1101/2025.03.25.645310

**Authors:** Pia Lavriha, Carina Fluri, Jorge Enrique Hernández González, Volodymyr M. Korkhov

## Abstract

Connexin gap junction channels (GJCs) are a family of proteins that connect the cytoplasms of two neighbouring cells and allow direct intercellular exchange of solutes smaller than ∼1 kDa, ensuring metabolic and electrical coupling of tissues. Connexin-32 (Cx32) GJCs mediate intercellular coupling in a variety of tissues, and specifically in the myelinating Schwann cells. Mutations in Cx32, such as W3S, are associated with peripheral neuropathy, X-linked Charcot-Marie-Tooth (CMT1X) disease. Phospholipids and sterols have been shown to regulate permeation through Cx32 GJCs, although the mechanism of regulation of Cx32 and other GJCs by lipids remains poorly understood. Here, we determine the cryo-EM structures of Cx32 GJCs reconstituted in nanodiscs, revealing that phospholipids block the pore of Cx32 GJC by binding to the site formed by N-terminal gating helices. The phospholipid-bound state is contingent on the presence of a sterol molecule bound to a hydrophobic pocket formed by the N-terminus: the N-terminal helix of Cx32 fails to sustain a phospholipid binding site in the absence of cholesterol hemisuccinate. Consistent with this, the CMT1X-linked W3S mutant which has an impaired sterol binding site adopts a conformation of the N-terminus incompatible with phospholipid binding. Our results indicate that different lipid species control connexin channel gating directly by influencing the conformation of the N-terminal gating helix.

**One-Sentence Summary:** Lipids modulate the gating helix conformation in connexin-32 channels

## Introduction

Cell communication is crucial for coordination of cells with their environment and generation of appropriate cellular responses. Among different means of cellular communication, gap junction intercellular communication (GJIC) enables a fast and direct signal exchange between neighboring cells and allows electrical and metabolic coupling of the tissues ^1,2^. In vertebrates GJIC is mediated by connexins, which assemble into gap junction channels (GJCs), that contain a central pore, connecting the cytoplasms of two neighbour cells and allowing solute permeation from one cell into the other ^1-3^. Each of the 21 connexin isoforms in humans has a distinct expression pattern, permeability, and gating properties, establishing the required GJIC between specific cell types ^4^. The GJIC has to be tightly regulated to ensure the passage of appropriate solutes and to prevent the exchange of noxious signals between coupled cells ^2-4^. Among the regulators, lipids (fatty acids, sterols, and phospholipids) have gained increased attention for their effects on GJC permeability. These hydrophobic small molecules have been shown to alter GJC permeability either directly by binding to the channel, or indirectly, via changes in the biophysical properties of the membrane or via activation of distinct signaling pathways ^5-8^.

The potential role of direct of lipid-protein interactions with the connexin channels has been illustrated by the recent structural studies of connexin hemichannels (HCs) and GJCs. Ordered lipid-like molecules have been identified at five different locations in recent cryo-electron microscopy (cryo-EM) structures of connexin HCs and GJCs: (i) as annular lipids, lining the extracellular lipid leaflet side of the channel (Cx46/50 GJC ^9^, Cx43 GJC ^10,11^, Cx36 GJC ^12^, and Cx32 HC and GJC ^13^), (ii) below the N-terminal helix (NTH) (Cx43 GJC ^10,11^, Cx43 HC ^11^, Cx32 HC ^13^), (iii) between NTHs (Cx43 GJC ^10,11^), (iv) lining the cytoplasmic entry of the pore (Cx36 GJC ^12^), and (v) lining the inside of channel pore (Cx31.3 HC ^14^, Cx36 ^12^, Cx46/50 ^9^, Cx32 GJC and HC ^13^ and Cx43 GJCs ^10,11^. The presence of these ordered lipid-like small molecules around or inside the channels suggests that lipids may have a direct effect on connexin channel structure and function.

Connexin-32 (Cx32) is a widely expressed isoform of the connexin channel family, found abundantly in the liver cells from which it was originally isolated ^15,16^. Cx32 is also found in the myelinating Schwann cells, where it appears to be important for maintaining the cellular homeostasis, with roles for both HCs and GJCs ^17^. Multiple mutations in Cx32 are associated with the X-linked Charcot-Marie-Tooth disease (CMT1X), a disorder accompanied by demyelination of the peripheral neurons and eventually leading to degeneration of the muscles, for which there is currently no cure ^17-20^. Cx32 HCs play a role in secreting ATP at the Schwann cell surface ^21^, and the GJCs are important to maintain lateral connections across the myelin layer ^22^. We have recently determined the first structures of human Cx32 HCs and GJCs in detergent micelles, where we observed a novel conformation of the N-terminal gating helix of Cx32 supported by a lipid-like molecule in the lipid-1 site, which we interpreted as a molecule of cholesterol hemiscuccinate (CHS) ^13^. Our work showed that two CMT1X-linked mutants, W3S and R22G, have a structural and a functional defect in the HCs, with little or no observable structural effects on Cx32 GJCs, and no measurable effects on Cx32-mediated GJC activity in cell-based assays ^13^.

In addition to our direct observation of sterol-like molecules bound to Cx32, the role of lipids in Cx32 regulation has been explored previously (**Figure 1A**). Cx32 preferentially resides in cholesterol-rich caveolin-containing lipid rafts ^8^. Phospholipids also directly associate with Cx32 and together with the sterols influence Cx32 HC permeability ^6^. However, how phospholipids and sterols interact with Cx32 channels in the context of a lipid bilayer remains unclear.

**Figure 1.**
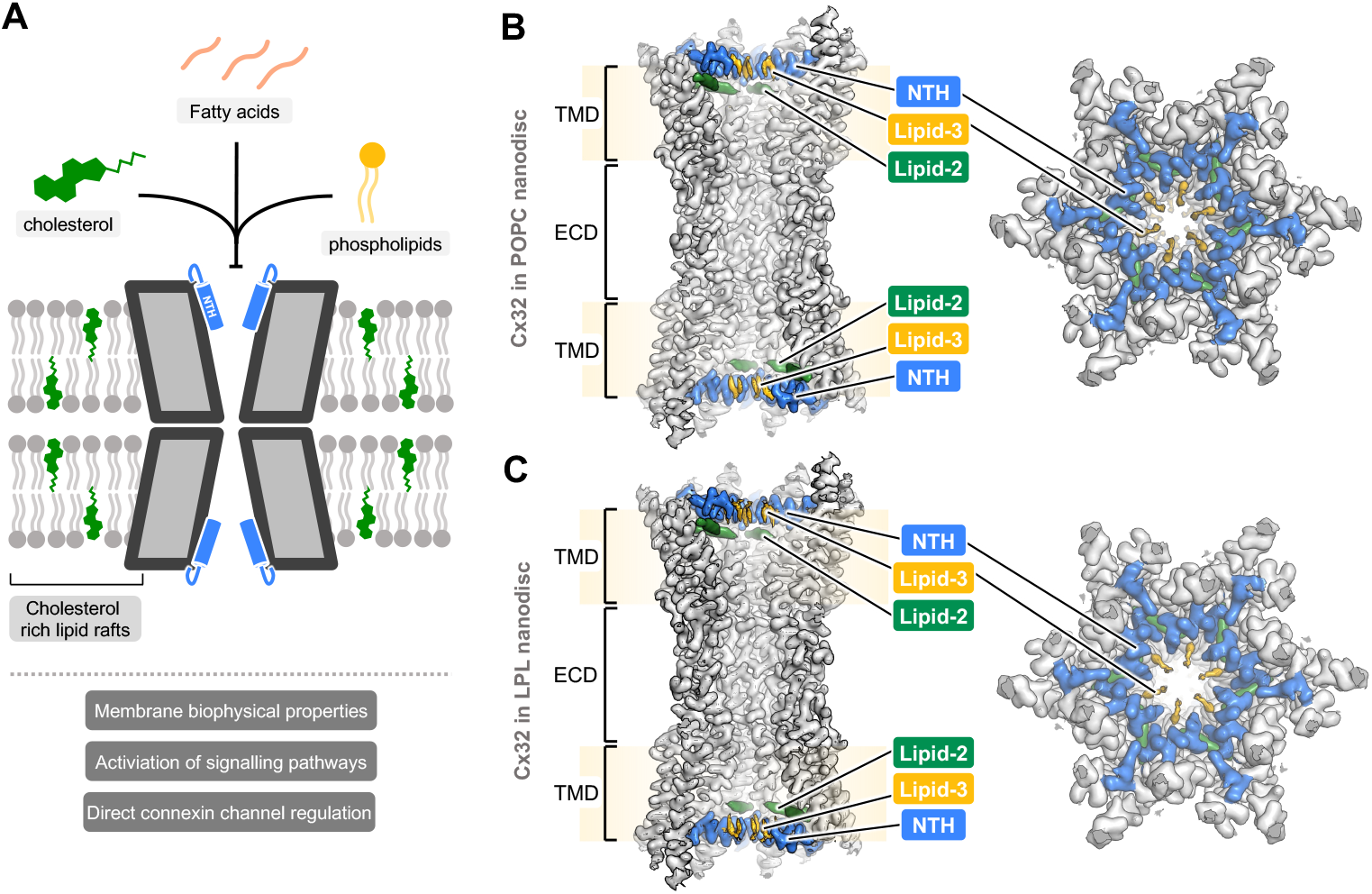
Cryo-EM structures of Cx32 GJC in nanodisc. **(A)** Schematic of the functional effects of lipids on Cx32 GJC. Structures of Cx32 GJC, reconstituted in nanodiscs, containing POPC **(B)** or LPL **(C)**, show an ordered N-terminal helix (NTH), and the presence of two densities, likely corresponding to lipids, lipid-2 and lipid-3. TMD – transmembrane domain, ECD – extracellular domain.

To investigate the effect of phospholipids and sterols on Cx32 channels, we reconstituted the purified Cx32 channels into lipidic environment in nanodiscs, and determined their cryo-EM structures.

## Results

### Structures of Cx32 GJCs in lipid environment

To address the effect of different lipids on Cx32 GJC, we purified Cx32 in digitonin, and reconstituted the protein into 1-palmitoyl-2-oleoyl-sn-*glycero*-3-phosphocholine (POPC) and liver polar lipids (LPL) -containing nanodiscs (Figure S1A-B). We chose two different lipid preparations for the following reasons: in the case of POPC we would obtain nanodiscs with clearly defined lipid content, and LPL would represent a close to physiological lipid composition, since Cx32 is known to be abundantly expressed in hepatocytes ^23^. We prepared samples for cryo-EM by plunge-freezing the purified and reconstituted samples in liquid ethane, collected the cryo-EM data, and determined the structures of Cx32 GJC in nanodiscs, at 3.20 Å and 3.29 Å resolution for POPC and LPL, respectively (Figures S2-3, S4A-B, Table S1). The structures were obtained by imposing the D6 symmetry. Refining the particles without imposing symmetry (C1) resulted in a similar reconstruction, albeit at a slightly lower resolution (Figures S2-3, S5-6).

Overall, the structures of Cx32 GJC in POPC- and LPL-containing nanodiscs are very similar to each other, with an RMSD of 0.495 Å (**Figure 1B-C**, Figure S6), and each was overall similar to the structure of Cx32 GJC in detergent ^13^. The four transmembrane helices (TM1-4) and the extracellular loops 1 and 2 (ECL1-2) of individual connexin subunits have the same conformation in the presence or absence of lipids, indicating that lipids do not affect the assembly of two HCs into a GJC, and do not affect the overall pore architecture. These similarities apply also to the orientation of the individual amino acid residues, with an RMSD of 0.259 Å and 0.636 Å between Cx32 GJC in detergent, and Cx32 GJC in POPC- and LPL-containing nanodiscs, respectively.

Nonetheless, the addition of lipids has a notable effect on the conformation of the Cx32 N-terminal helix (NTH), an *α*-helical region critical for Cx32 gating ^13,24,25^ (**Figures 2-4**). The NTH transitions from a disordered to an ordered α-helical conformation, as well as moves from lining the channel pore to pointing towards the symmetry axis of the pore. This movement is independent from conformational changes in the TM1-4 or ECL1-2. This conformation resembles the NTH conformation of Cx32 HC that we previously determined using Cx32 samples in detergent ^13^. Performing protomer focused classification (PFC) on the individual subunits of Cx32 GJC in nanodiscs did not expose any substantial variability in the NTH conformation (Figure S7A and B), indicating that the NTH of all subunits in these conditions assume a similar conformation.

**Figure 2.**
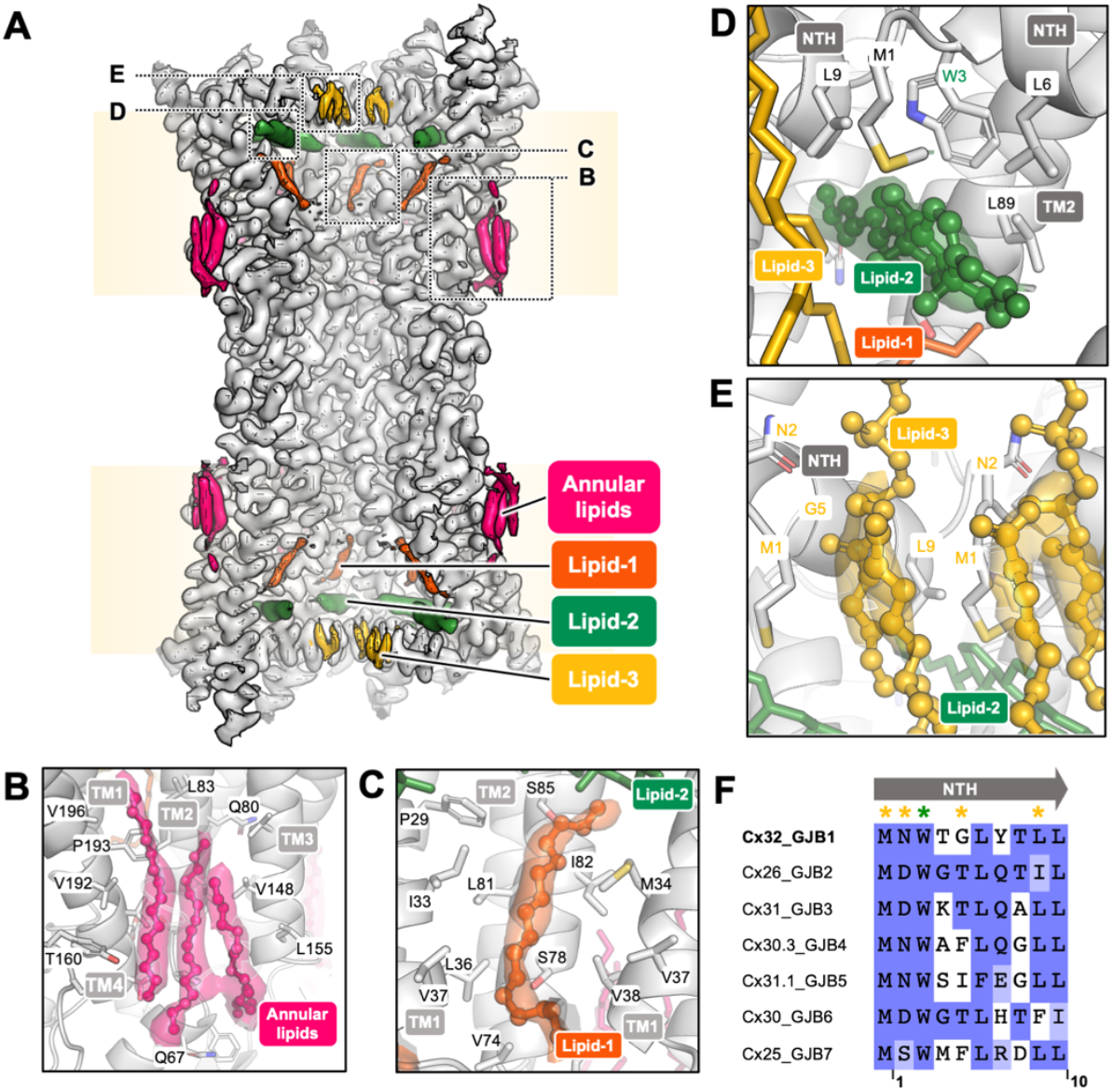
Lipids bind to Cx32 GJC in nanodisc. **(A)** Lipid densities in Cx32 GJC nanodisc structure (POPC-containing nanodiscs). Dashed boxes represent the positions of lipid densities, represented in panels B-E. The binding sites of annular lipids **(B)**, lipid-1 **(C)**, lipid-2 **(D)** and lipid-3 **(E). (F)** Multiple sequence alignment of β-group connexins, performed using Clustal Omega algorithm. The alignment is colored by BLOSUM62 color scheme. Yellow asterisks represent the putative lipid-3-interacting residues, and green asterisk indicates the W3 residue, interacting with lipid-2.

### Annular and pore-lining lipids

Several lipid-like densities have been identified in the Cx32 GJC structure in detergent. In particular, lipid-1, which lines the inside of the channel pore, and annular lipids on the extracellular membrane leaflet side of the channel. Due to the limited resolution of the 3D reconstructions, the lipid identities could not be unequivocally determined ^13^. The densities at equivalent positions in Cx32 GJC structure in nanodisc are resolved better and could correspond to acyl chains (**Figure 2A**). Acyl chains in similar positions were identified in the structures of Cx46/50 GJC ^9^, Cx43 GJC ^10^, and Cx36 GJC ^12^. As we were not able to determine whether these acyl chains represent free fatty acids copurified with the protein, or whether they are parts of bound phospholipids, we left these regions of the EM maps unmodelled (the hexadecane in **Figure 2** is shown for illustration purposes).

Annular lipids are present at the interface between two neighbouring connexin subunits (**Figure 2B**). The lipid binding pocket is formed by TM1 and TM4 of one subunit and TM2 and TM3 of the neighboring subunit. The pocket is formed predominantly by hydrophobic amino acids. Similarly, lipid-1 binds to a hydrophobic pocket formed by neighboring connexin subunits, in this case formed by TM1 of one and TM1 and TM2 of the second (**Figure 2C**). The lipid-1 density spans the majority of the transmembrane domain, from lipid-2 below the NTH to the interface between the transmembrane and extracellular domains.

### Sterol-like and phospholipid-like molecules stabilize the NTH

Closer inspection of the density map around the NTH in both Cx32 GJC nanodisc structures, reveals two densities which could stabilize the observed NTH conformation (**Figure 2A**). The first is positioned below the NTH (**Figure 2D**) and is shaped like a sterol. This density is similar to the one we have observed at the lipid-2 site previously for Cx32 HCs in detergent ^13^, as well as in the structures of Cx43 GJCs solved in both nanodiscs and in detergent ^10,11^. However, the identity of the lipid-2 molecule could not be determined definitively. Although we modelled a cholesterol molecule into the corresponding part of the density map, this density could also accommodate other sterols, such as cholesteryl hemisuccinate (CHS) and digitonin, which were used during stages of protein purification, as described in Materials and Methods (Figure S8A). This binding site is hydrophobic, with the W3 residue side chain in direct proximity, participating in the interactions with the bound lipid molecule. Residue W3 is conserved in all β-group connexins (**Figure 2F**), suggesting its functional importance in interactions with the lipid-2 as a mediator of NTH conformational changes in Cx32 and other GJCs.

An additional observed density, which we named lipid-3, is located between the neighboring NTH regions. The lipid-3 density has a similar shape in both POPC- and LPL-containing Cx32 GJC nanodisc structures (**Figure 2E**, Figure S8B,C). The density is consistent with the glycerophosphoric segment and for parts of the two aliphatic chains of POPC. The binding site for lipid-3 is formed by residues M1, N2, G5 and L9 of one Cx32 subunit and M1 of the neighboring subunit. The position equivalent to G5 in other β-group connexin isoforms is not conserved but occupied by hydrophobic residues (**Figure 2F**), which could mediate the lipid-3 interaction in other β group connexins. Mutation of one of the conserved residues, L9W, is associated with CMT1X ^26^, indicating the functional importance of this site.

### Cx32 reconstituted in nanodiscs is effectively plugged by lipid molecules

The structure of Cx32 GJC reconstituted into nanodiscs shows a conformational change of the NTH that leads to a decrease in the pore diameter from approximately 15 Å in an open conformation to 12 Å (**Figure 3, A and B**, Figure S8D). This NTH reorganization modifies the electrostatic potential of the cytoplasmic pore entry, which becomes more positively charged (**Figure 3C**). We refer to this new Cx32 GJC conformation as the ordered NTH state (NO-state). Including lipid-3 in pore diameter calculations shows an additional restriction of the pore diameter to only 4 Å (Fig**ure 3, A and B**), which effectively constricts and blocks the pore entry to ions or small molecules.

**Figure 3.**
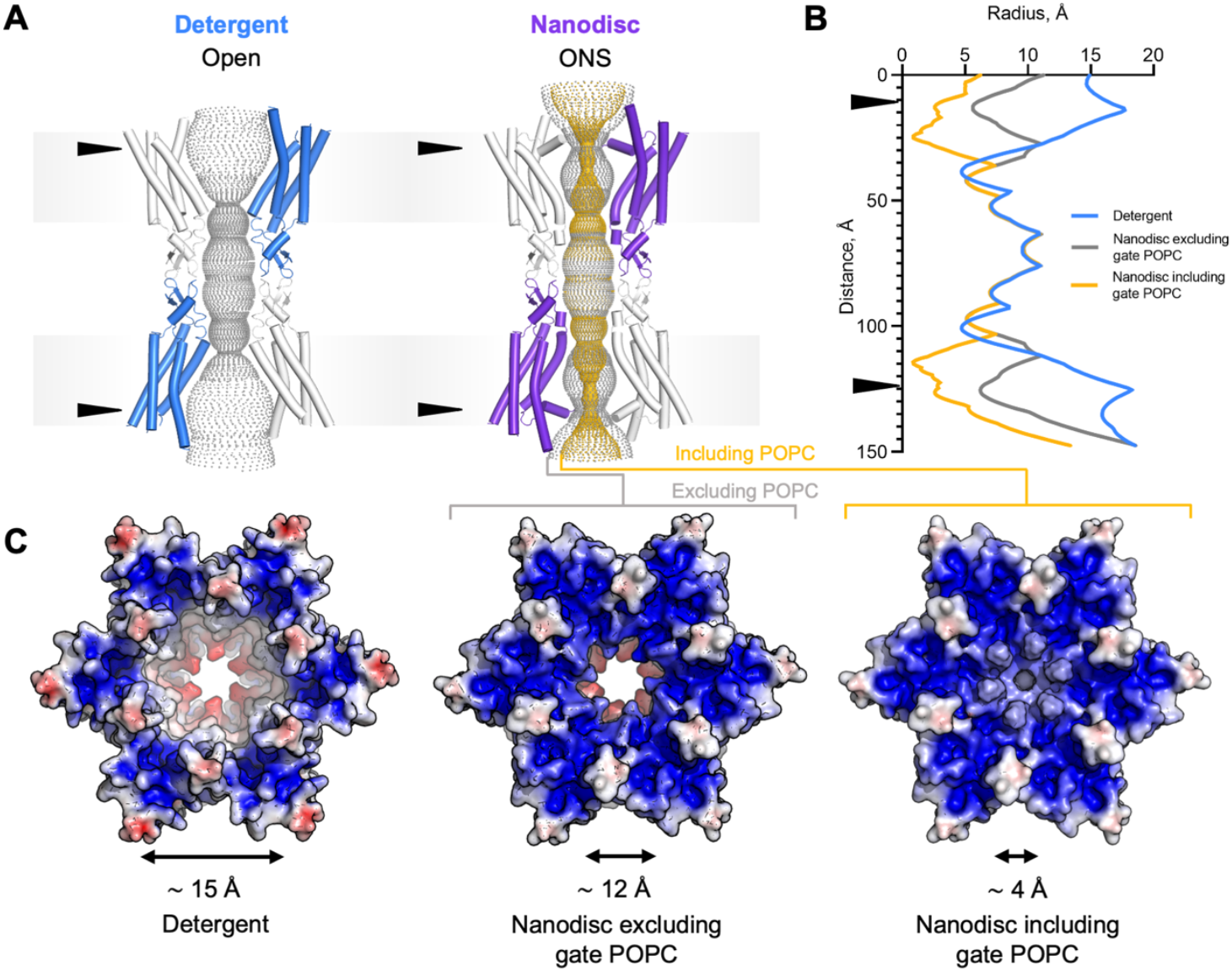
Lipids induce changes in Cx32 GJC surface electrostatic potential and pore diameter. **(A)** Pore representations of Cx32 GJC solved in detergent and Cx32 GJC in nanodisc, calculated using the HOLE method. Gray contour represents the pore diameter without and yellow with POPC in the HOLE calculation. The arrows indicate the position of change in the pore radius due to NTH rearrangement.NO-state – ordered N-terminus state **(B)** The pore radius of Cx32 GJC, represented in **(A)**, calculated using HOLE. Both nanodisc plots represent the POPC nanodisc conditions, including or excluding POPC in the calculation. **(C)** Electrostatic surface potential of Cx32 GJC, solved in detergent, in nanodisc, and in nanodisc including POPC in electrostatic surface calculation.

### Lipid-2 binding is required for NTH stabilization

The positions of lipid-2 and lipid-3 in the vicinity of the Cx32 NTH regions, known to be the key gating regions of connexin channels, suggest that these two lipids may stabilize NTH in the NO-state. To assess the requirement for lipid-2 and lipid-3 in NTH stabilization, we determined the GJC structures of (i) Cx32 GJC purified without addition of the cholesterol analog CHS during protein purification, and (ii) Cx32 W3S mutant, which has an impaired lipid-2 binding site ^13^ (Figures S1C-D, S4, S9-S11). The structures were determined at 3.12 Å and 2.35 Å resolution, respectively, by imposing D6 symmetry, although the same reconstruction at a slightly lower resolution was obtained in C1 symmetry (Figure S5).

The Cx32 GJC structure in nanodisc without addition of CHS during purification confirmed that the density corresponding to lipid-2 is indeed CHS, and this molecule is a prerequisite for stabilizing the NO-state (**Figure 4A-B**). Removal of CHS from the purification procedure, followed by nanodisc reconstitution of Cx32, results in a 3D reconstruction where NTH tilts and aligns along the channel pore, displacing the lipid-2 density (**Figure 4B**). This conformation is compatible with an open state of a GJC, similar to those observed for Cx26 ^27^ (**Figure 4D**, Figure S12).

**Figure 4.**
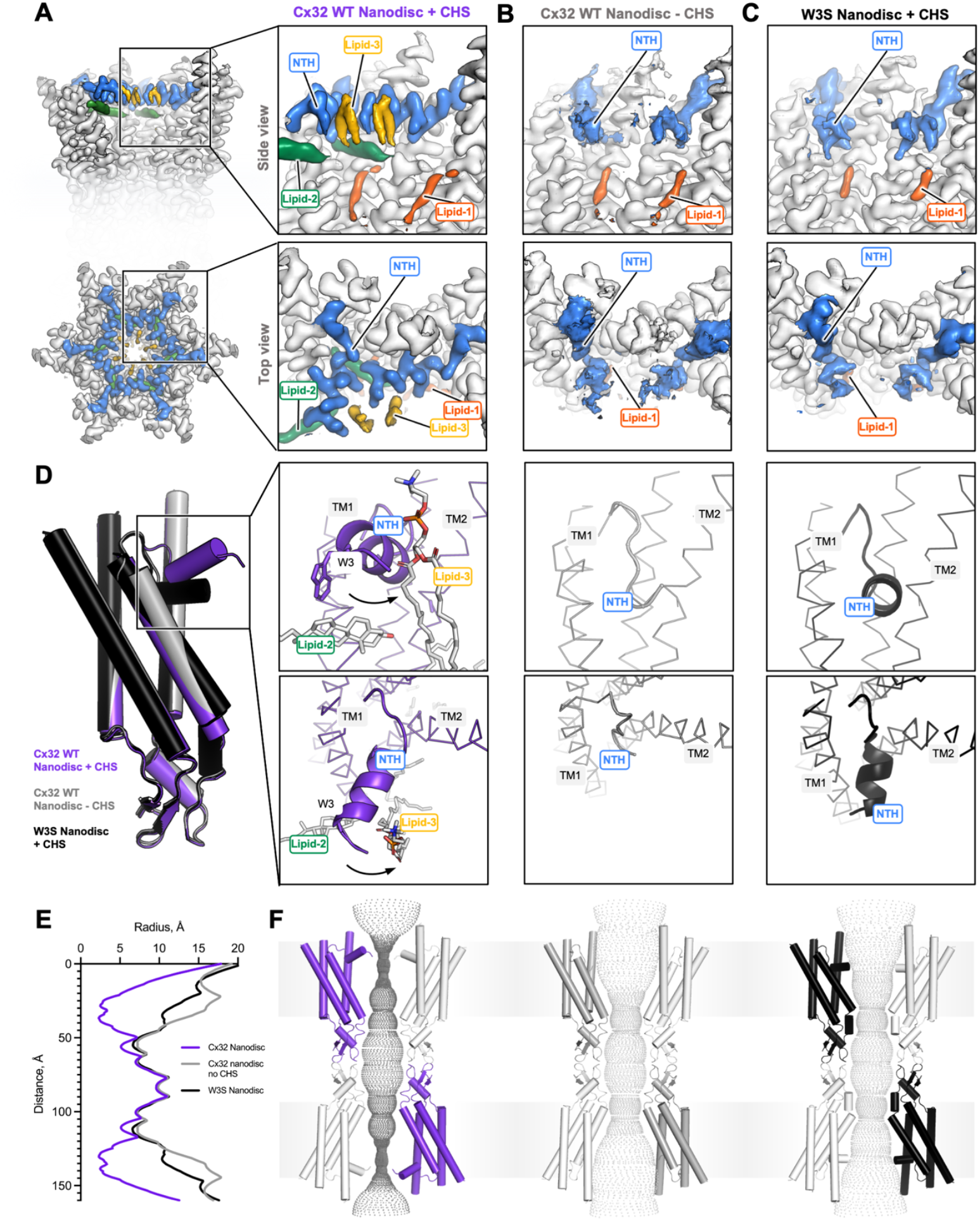
Lipid-2 is necessary for NTH stabilization in NO-state. **(A)** Presence of both lipid-2 and lipid-3 results in an ordered N-terminal helix (NO) confirmation. **(B)** Removing CHS from Cx32 reconstitution into nanodisc results in a different conformation of the NTH, which appears to be less stable and does not contain a lipid-2 nor lipid-3 density. **(C)** W3S mutant, which has an impaired lipid-2 binding site, similarly does not have a lipid-2 nor lipid-3 density and contains a less ordered and different NTH conformation from NO-state. **(D)** The absence of lipid-2 leads to NTH rearrangements, which are different from the NO-state state (Cx32 WT Nanodisc + CHS). **(E)** Graphical representation of pore diameter analysis performed using HOLE, represented in (F). **(F)** Pore diameter analysis of NO-state (Cx32 Nanodisc + CHS), Cx32 WT Nanodisc - CHS, and W3S Nanodisc + CHS, calculated using HOLE software.

### W3S GJCs do not form an NO-state in lipidic environment

Mutation of the residue W3 to a serine produced a similar effect on Cx32 GJCs upon lipid reconstitution in the presence of CHS, as removing CHS did on the wild-type Cx32 (**Figure 4C**, Figure S13A). The NTH of Cx32 GJC in POPC is ordered and points towards the inside of the channel pore, in a conformation slightly different from that in the wild-type Cx32 maintaining a pore diameter of ∼10 Å (**Figure 4D-F**, S12). The conformation of the NTH in the W3S GJC in lipidic environment is distinct from the constricted NTH conformation observed in the Cx32-W3S HC (Figure S13B) ^13^.

To determine whether there is a variability in the NTH conformation among the individual subunits of the W3S and the wild-type Cx32 (in the absence of CHS) GJCs, and whether any of the Cx32 subunits still feature lipid-2 or lipid-3 densities that may be averaged out due by imposing the D6 symmetry, we performed protomer-focused classification (Figure S7C-D). In all classes for both analysed datasets we could observe the NTH in only one corresponding major conformation. Neither lipid-2 nor lipid-3 could be identified in the classes, further confirming that the lipid-2 corresponds to the CHS molecules added during purification.

Altogether, our results with the wild-type Cx32 in the absence of CHS and the Cx32-W3S GJCs indicate the requirement of the sterol binding site (lipid-2 site) for stabilizing the NTH in a conformation that allows the phospholipids to bind to the lipid-3 sites.

### Analysis of Cx32 NTH motions through molecular dynamics simulations

To assess the dynamic interplay between the NTH and the lipids present in the pore of Cx32, we performed molecular dynamics simulations (MD) using a minimal hexameric Cx32 HC system, embedded in a lipid bilayer (see Materials and Methods for details on the system setup).

For the purposes of our MD analysis, we used cholesterol (CHOL) instead of CHS, because CHOL is a natural sterol that is abundant in biological membranes and is likely to occupy the lipid-2 binding sites in the native Cx32 GJCs / HC. Secondary structure analysis was performed for NTH residues 3-10 of each chain in wild-type Cx32 and Cx32 W3S HCs both in the presence and absence of bound lipids, POPC and CHOL. We monitored the average probability of the aforementioned residues adopting either an α-helix or random coil conformation over the course of two replicate 1 μs MD simulations for each studied system (**Figure 5A-B, E-F**). The results indicate a higher probability of residues 3-10 adopting a random coil conformation in Cx32 W3S compared to wild-type Cx32 (**Figure 5A-B**). Interestingly, the disruption of the α-helix secondary structure becomes more significant in Cx32 W3S as the simulation time increases (**Figure 5B**). This underscores the instability of N-terminal α-helices carrying the W3S mutation, initially modeled as in the native protein. On the other hand, the presence of bound POPC and CHOL was found to enhance the stability of the NTHs in both wild-type Cx32 (**Figure 5E**) and Cx32 W3S (**Figure 5F**) relative to the respective apo forms (**Figure 5A-B**).

**Figure 5.**
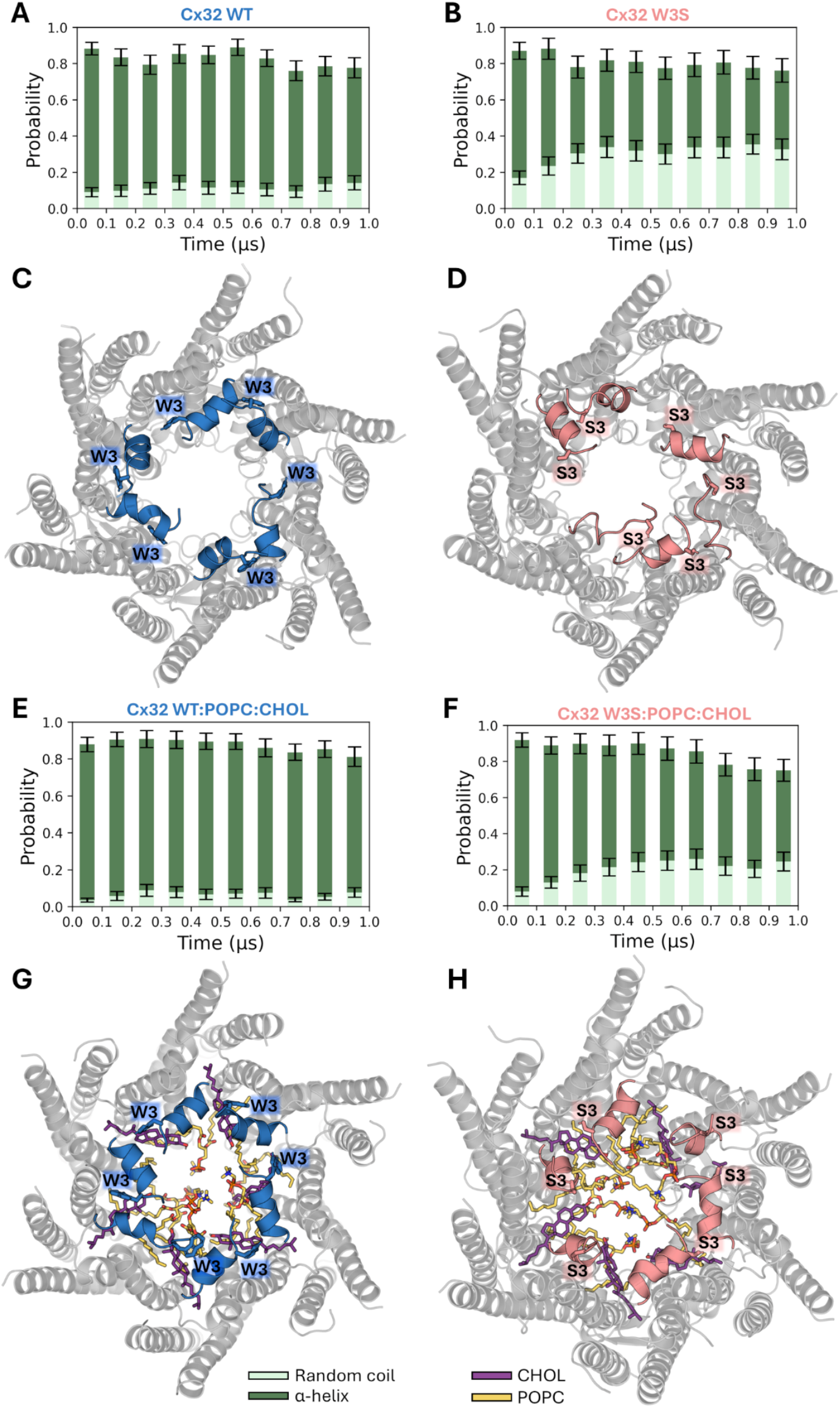
Time evolution of the secondary structure of residues 3-10 in different Cx32 systems during 1 μs MD simulations. **(A-B), (E-F)** represent the mean probabilities of residues 3-10 (which form the N-terminal α-helices in Cx32 wt) adopting an α-helix or random coil conformation for the indicated systems throughout the simulation time. For each system, the plotted mean probabilities were calculated by averaging the per-residue probabilities for residues within the 3-10 interval in every protein chain and over the two replicate MD simulations in 0.1 μs intervals (i.e., 0-0.1, …, 0.9-1 μs). Other types of secondary structure are omitted. (**C-D), (G-H)** show a structural representation of the last frame (*t=*1 μs) in one of the two replicate trajectories for each system. The N-terminal residues 1-10 are colored in blue and salmon on each panel.

Structural representations of the final frames collected for each analyzed system in one of the replicate 1 μs trajectories are provided to visualize the conformational changes undergone by the NTH residues during the MD simulations (**Figure 5C-D, G-H**). The depicted structure of apo Cx32 W3S hexamer (**Figure 5D**) clearly illustrates the tendency of some of its NTHs to unfold during the microsecond-long MD simulation. These results suggest that longer MD simulations may be required to observe the unfolding process of all the six NTHs. To a lesser extent, a similar phenomenon can be observed in the last frame of Cx32 W3S in the presence of bound lipids (**Figure 5H**). In contrast, the N-terminal α-helices of wild-type Cx32 remained largely intact at the end of the MD simulations in both conditions (**Figures 5A** and **5G**). The conclusions drawn from the visual inspection of the last frames are consistent with the secondary structure analysis presented above.

We extended the same analysis to four systems consisting of wild-type Cx32 and Cx32 W3S, each bound to either CHOL or POPC (Figure S14). For these new systems, we confirmed the higher propensity of the Cx32 W3S NTHs to unfold compared to the native protein in the same conditions. Furthermore, the presence of either CHOL or POPC stabilized the N-terminal α-helices in both wild-type Cx32 and Cx32 W3S relative to their apo forms.

The RMSF and RMSD calculations for the NTHs (Figures S15 and S16) underscore the increased flexibility in the N-terminal residues of Cx32 W3S mutant across all conditions, becoming more significant in the absence of lipids. Additionally, RMSD measurements reveal a greater tendency of the N-terminal residues in the mutated protein to deviate from their initial α-helix conformation compared to those in wild-type Cx32.

The influence of lipids on NTH motion relative to TM1 was investigated by monitoring an angle during MD simulations, defined by the centers of mass of three atom groups at the NTH, the N-terminal end of TM1, and the center of TM1 (see Figure S17). The angle distributions for wild-type Cx32 show that neighboring NTHs tend to alternate between conformations shifted upward and downward relative to the position observed in the cryo-EM structure of Cx32 in nanodiscs (Figure S18). A similar trend was observed for Cx32 W3S, with broader angle distributions reflecting partial NTH unfolding (Figure S19). In contrast, for wild-type Cx32 and W3S with POPC and CHOL at the lipid binding sites, the angle distributions are narrower, suggesting that the bound lipids restrict the NTH motion to positions closer to that observed in the experimental structure (Figures S20 and S21).

Overall, the results presented here highlight the destabilizing effect of the W3S mutation on the Cx32 NTH. Moreover, we observed that the binding of CHOL and/or POPC exerts a stabilizing influence on these helices in both the mutant and wild-type proteins. Disruption of the CHOL binding site due to the unfolding of the NTH regions in the W3S mutant makes the presence of ordered lipids within the pore less likely based on our cryo-EM data.

### Dynamic behaviour of lipids within the Cx32 pore region

To investigate the conformational dynamics of lipids bound to wild-type Cx32 and Cx32 W3S during the 1-μs replicate MD simulations, we tracked the RMSD values for their heavy atoms relative to the corresponding starting conformations in the initial structure (**Figure 6A-H**). The distribution of RMSD values for CHOL indicates that these molecules remained largely in native-like conformations when bound to the wild-type protein, as evidenced by a high and narrow peak around ∼2.5 Å (**Figure 6B**). In contrast, a wider RMSD distribution was observed for CHOL bound to Cx32 W3S (**Figure 6B**), likely due to partial disruption of the NTH residues in the mutated system (**Figures 5F** and **5H**). This structural change may compromise the integrity of the CHOL binding site.

**Figure 6.**
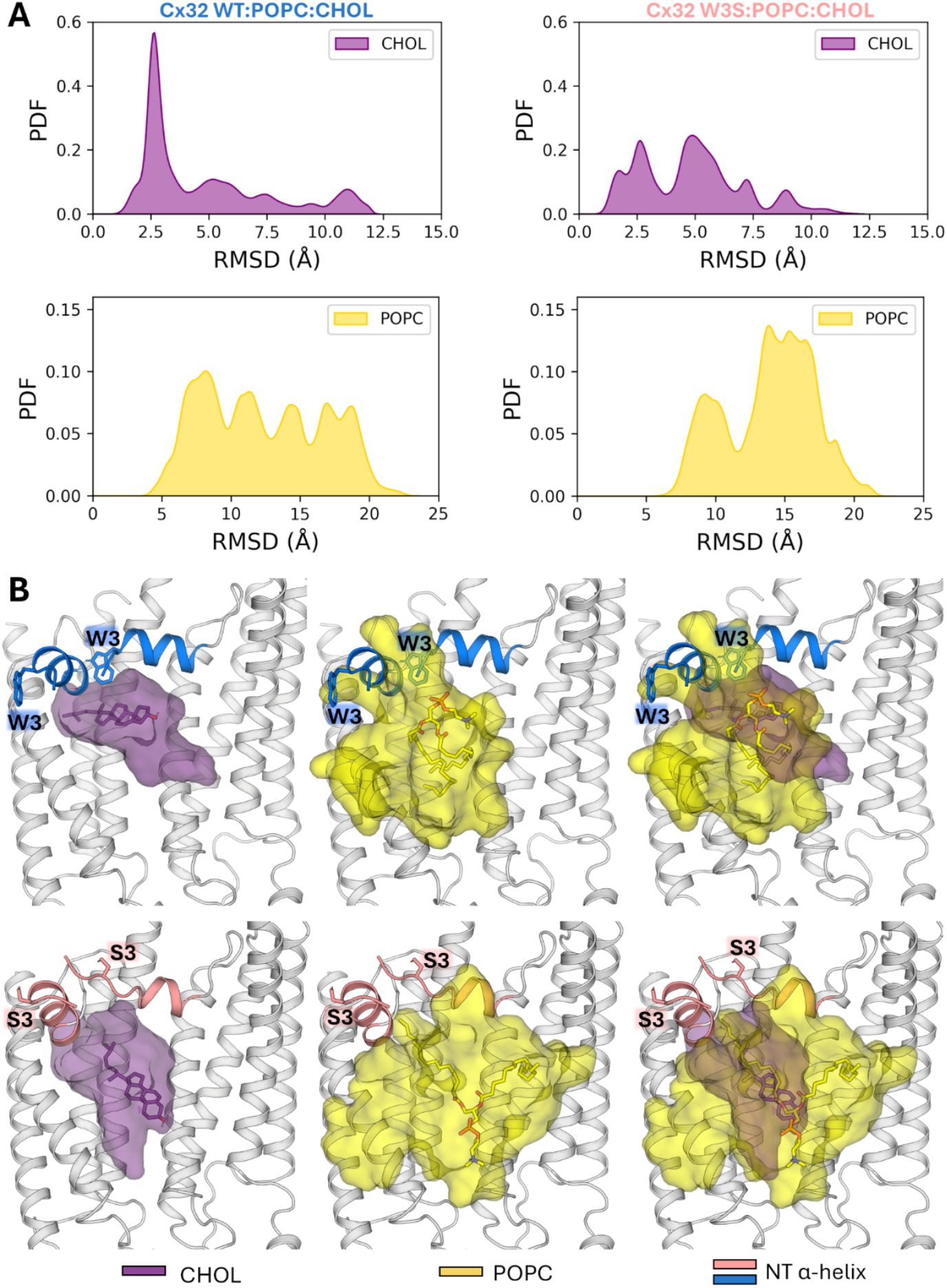
Dynamic behavior of lipids within the Cx32 pore region during MD simulations. **(A)** Distribution of RMSD values for the heavy atoms of the six CHOL molecules bound to Cx32 in the Cx32 wt:POPC:CHOL and Cx32 W3S:POPC:CHOL complexes. RMSD values were calculated relative to the minimized initial structure, with all trajectory frames from two 1-μs replicate simulations for each system fitted to the protein backbone in the reference structure. The first 100 ns of each trajectory was discarded to ensure equilibrium in the distributions. **(B)** Structural depictions of the central structures for Cx32 wt:CHOL:POPC and Cx32 W3S:CHOL:POPC, calculated through clustering analysis. Transparent surfaces enclosing the POPC and CHOL molecules were generated after superimposing all protein chains along with the bound lipids within each system. One CHOL molecule and one POPC molecule, located at the centers of their respective accessible regions, are depicted as sticks, while two adjacent protein chains forming the complete CHOL binding site are shown as cartoons. For clarity, three panels are presented for each system: one depicting each lipid individually and one showing both lipids together.

In both systems, POPC molecules exhibit broad RMSD distributions throughout the simulations, reflecting their high intrinsic flexibility and binding to a relatively shallow surface on the protein (**Figure 6A**). Notably, the main peak of RMSD values for the POPC molecules bound to Cx32 W3S is shifted toward higher values compared to the wild-type system’s distribution. This suggests that the mutation can also compromise the binding of POPC to the NTH region of Cx32.

**Figure 6B** shows the central structures of Cx32 wt and Cx32 W3S bound to POPC and CHOL, determined through clustering analysis. Transparent surfaces were generated to enclose the conformational space explored by the ligands’ central structures after superposition of all chains within each system. These structures highlight the loss of the native binding mode of CHOL to Cx32 W3S and reveal a tendency for POPC molecules to adopt conformations in the mutant that are more detached from the NTH compared to the wild-type system. Thus, the MD simulations suggest that CHOL and POPC exhibit stronger binding to wild-type Cx32 than to Cx32 W3S, consistent with experimental findings.

## Discussion

The structures of Cx32 GJC in different lipidic environments provide a mechanistic insight to the previously suggested effects of lipid composition on Cx32 channel function ^5-7^ and NTH-mediated gating ^13^. The NTH is a dynamic element of the connexin channel structure, reacting to the presence of different lipid species by changing its conformation and thus regulating the solute permeation through the Cx32 GJCs (**Figure 7**). In the presence of phospholipids (POPC or LPL) the NTH reorient from lining the channel pore (observed in the open conformation) to pointing toward the channel pore axis. However, binding of sterols to the lipid-2 site is required for this. The switch from open to NO-state is associated with changes in both the electrostatic properties in the radius of the pore, suggesting that these NTH transitions change the range of molecules that can pass through the channel.

**Figure 7.**
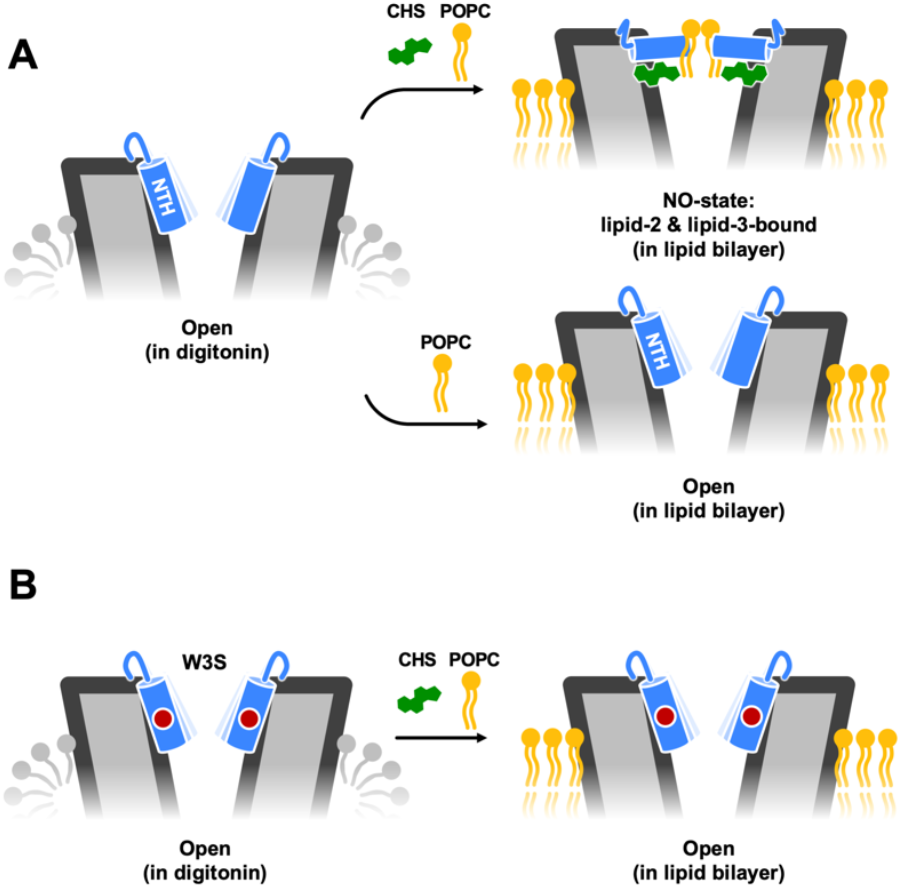
A schematic representation of the N-terminus changes in Cx32 GJC in response to changes in lipid composition. **(A)** Structures of Cx32 GJCs determined using protein in detergent micelles feature a dramatically different conformation of the NTH, compared to those determined in lipid bilayer (in nanodiscs). The observed conformations depend critically on the presence of sterols (CHS). The NTH is represented in blue, the sterol molecules in green, and phospholipids in yellow. **(B)** The CMT1X disease-linked mutation W3S is insensitive to the sterol effects on the NTH structure. These cryo-EM observations are consistent with the MD simulations, which reveal a mutually stabilizing role of the lipids and the NTH residues.

It is important to note that our MD simulations reveal a layer of complexity in lipid-protein interactions that is not apparent when observing the “static” cryo-EM structures. Our cryo-EM reconstruction of the Cx32 GJC in lipidic environment indicates that a conformation of the NTH is formed whereby phospholipids and sterols work together to lock the protein in a particular conformation. In stark contrast, our MD simulations reveal a great degree of plasticity of the lipid binding sites within the pore, especially the putative phospholipid interactions sites formed by the NTH regions, as suggested by the cryo-EM reconstructions. It remains to be determined whether additional factors play a role in stabilizing both the lipids and the Cx32 NTH residues.

We have observed previously that the CMT1X-causing Cx32 mutant W3S has a closed NTH conformation in the cryo-EM structures of W3S HC in detergent, correlating with the functional effect on the W3S HC function ^13^. Interestingly, the Cx32 GJCs appeared unaffected by the W3S mutation. The results presented here suggest that reconstituting Cx32 into a lipidic environment allows us to sample the phospholipid-induced conformational changes of the NTH: while wild-type Cx32 is capable of interacting with POPC (or other phospholipids present in the LPL mixture) via the lipid-3 site in the lipid-2-dependent manner, the W3S mutant fails to establish the lipid-2 site and thus does not form a lipid-3-bound NO-state.

Previous structural studies on Cx43 GJC ^10,11^, Cx36 GJC ^12^, and Cx32 HC ^13^ have implicated the NTH conformational changes in regulation of the connexin channel permeability and suggested a link between these conformational changes and distinct lipid environments. However, it remains to be determined whether phospholipids and sterols bind and induce the observed NTH rearrangements in a physiological context. It is tempting to suggest that localization of Cx32 in cholesterol-rich membrane microdomains ^8^ may predispose Cx32 to interactions with cholesterol via various sites at the outer or inner surface of the channel. However, the path taken by either cholesterol or the phospholipids to reach the lipid-2 or lipid-3 sites, respectively, is not immediately obvious, as these discrete sites are separated from the lipid bilayer by the protein. Delineation of the route used by the lipids to dynamically regulate Cx32 channels via lipid-2 and lipid-3 sites will require careful experimental and computational analysis.

It is worth noting that gating by lipids has become a topic of intensive investigations not only in connexins, but also in other members of the large pore channel family. For example, obstruction of the pore by phospholipids has been observed in the *Caenorhabditis elegans* innexin-6 channels reconstituted into nanodiscs ^28^. The volume-regulated LRRC8A:C channels have been shown to utilize the intra-pore lipids to block ion conduction ^29^. Another member of the family, the calcium homeostasis modulator 1, CALHM1, has been shown to accommodate sterols and phospholipids in a conserved lipid-binding pocket; in the case of CALHM1 phospholipid binding appears to stabilize and regulate the channel ^30,31^. Lipid-mediated regulation is a common denominator in these studies, and clear parallels can be drawn with our own observations on lipid-reconstituted Cx32 channels.

## Acknowledgments

JEHG thanks the National Laboratory for Scientific Computing (LNCC/MCTI, Brazil) for providing HPC resources of the SDumont supercomputer, URL: http://sdumont.lncc.br. We thank the PSI scientific computing team for expert support in high performance computing and image analysis. JEHG thanks the financial support of the National Council for Scientific and Technological Development (CNPq), Grant 153794/2024-0, and of the Sao Paulo Research Foundation (FAPESP), Grants 2020/08615-8, 2022/03901-8, 2024/13327-2. The work was supported by a Swiss National Science Foundation to VMK (184951).

## Author contributions

Conceptualization: VMK

Methodology: PL, JEHG, VMK

Investigation: PL, CF, JEHG, VMK

Visualization: PL, JEHG, VMK

Funding acquisition: VMK

Project administration: VMK

Supervision: VMK

Writing – original draft: PL, JEHG, VMK

Writing – review & editing: PL, JEHG, VMK

## Data and materials availability

The MD simulation trajectories and data have been deposited to the OSF repository (https://osf.io/gezpx/). All coordinates and cryo-EM density maps have been deposited to the Protein Data Bank (9QN9, 9QND, 9QNF, 9QNT) and Electron Microscopy Data Bank (EMD-53240, EMD-53244, EMD-53245, EMD-53250).

## Supplementary Materials

Materials and Methods

Fig. S1 to S21

Table S1

References (1-18)

