## Supplementary Materials for "Lipid dependence of connexin-32 gap junction channel conformations"

### Materials and Methods

**Lipid Preparation.** The lipids (1-palmitoyl-2-oleoyl-glycero-3-phosphocholine (POPC) or bovine liver polar lipid extract (LPL)) dissolved in chloroform at a concentration of 25 mg/ml were transferred to a glass tube and dried under the nitrogen stream until completely dried. A buffer consisting of 25 mM Tris-HCl pH 8.0, 150 mM NaCl, 2 % n-Dodecyl β-D-maltoside (DDM) was added to reach lipid concentration of 8.33 mg/ml. The lipids were placed in a sonication bath at 45 °C until the lipids were completely dissolved; the clarified lipid stocks were aliquoted and stored at - 20 °C until use.

**MSP2N2 expression and purification.** A single colony of *E. coli* BL21(DE3) cells transformed with MSP2N2 plasmid was used to inoculate 100 ml of LB medium supplemented with 50 µg/ml kanamycin and grown over night at 37 °C and 180 rpm. Terrific broth (TB) medium (1L), supplemented with 50 µg/ml kanamycin, was inoculated with 30 ml of the overnight culture, and grown at 37 °C and 180 rpm to the $OD_{600}$ of ~3.0. Protein expression was induced by addition of IPTG at a final concentration of 1 mM for 3 h at 37 °C and 180 rpm. Cells were harvested by centrifugation for 30 min at 4000 rpm and 4 °C. The cell pellets were resuspended in 50 ml of 40 mM Tris-HCl pH 8.0, 0.3 M NaCl and 1 % Triton X-100 per 1L of cell culture, supplemented with 1 mM PMSF and 10 µg/ml DNase I and lysed by sonication for 15 min on ice at 50 % amplitude. The lysate was clarified by centrifugation for 30 min at 20000 x g. The cleared lysate was added to Ni-NTA resin (2 ml per 1 L of cell culture) and incubated for 30 min at 3 °C with constant rotation. The resin was collected in a gravity column, washed with 20 column volumes (CV) of 40 mM Tris-HCl pH 8.0, 0.3 M NaCl, 50 mM Na-cholate and 20 mM imidazole, then 20 CV of the same buffer with 50 mM imidazole and eluted with 5 CV of the same buffer with 400 mM imidazole. The protein was desalted into 20 mM Tris-HCl pH 8.0, 0.1 M NaCl, 0.5 mM EDTA using G-25 PD-10 desalting column, concentrated using an Amicon Ultra 10 kDa cut-off concentrator, aliquoted and stored at -80 °C until use.

**Cell Culture.** HEK293F cells were maintained in 15 cm plates in Dulbecco's Modified Eagle Medium (DMEM) supplemented with 10 % Fetal Calf Serum (FCS) and 1 % Penicillin-Streptomycin (PenStrep) at 37 °C and 5 % $CO_2$. For transfection using branched polyethyleneimine (PEI), the medium was replaced with DMEM supplemented with 2 % FSC PenStrep. The DNA and PEI dilutions were prepared separately in un-supplemented DMEM, using 40 µg DNA per plate and PEI in 1:2 ratio of DNA to PEI (w/w). The dilutions were mixed and incubated at room temperature for 5 min and added to the cells in the drop-wise manner. The cells were cultured at 37 °C and 5 % $CO_2$ for 48 h, after which they were harvested by scraping and stored at -80 °C.

**Connexin Purification.** The protein purification was performed as described previously [1]. The cells harvested from 100 15-cm culture plates were resuspended in buffer A (25 mM Tris-HCl pH 8.0, 150 mM NaCl) supplemented with protease inhibitors (1 mM benzamidine, 1 µg/ml leupeptin, 1 µg/ml aprotinin, 1 µg/ml pepstatin, 1 µg/ml trypsin inhibitor and 1 mM PMSF) and 10 µg/ml DNase I. The cells were lysed by 300 sonication pulses at 35 % amplitude (Sonics Vibra-Cell, cycle: 0.5 s pulse on, 0.5 s pulse off). The lysate was clarified by ultracentrifugation at 4 °C, 35 000 rpm (Ti45 rotor, Beckmann Coulter) for 50 min. The pellet was transferred to fresh buffer A supplemented with protease inhibitors and homogenized. The membranes were solubilized by addition of 1 % DDM and 0.2 % cholesteryl hemisuccinate (CHS) for 1 h at 4 °C, with constant rotation. In the case of purifying Cx32 in the absence of CHS, CHS was omitted in this membrane solubilization step. The membranes were clarified by ultracentrifugation at 4 °C, 35 000 rpm (Ti45 rotor) for 50 min. The solubilized and clarified membranes were incubated with 2 ml of anti-GFP nanobody coupled CNBr-sepharose resin for 30 min at 4°C, with constant rotation. The resin was applied to a gravity column, washed with 40 column volumes of buffer C (25 mM Tris-HCl pH 8.0, 150 mM NaCl, 0.1 % digitonin). The protein was eluted by cleavage using 3C protease (0.4 mg) for 2 h with rotation at 4 °C. The eluted protein was concentrated to a volume <800 µl using AmiconUltra 100 kDa cut-off concentrator.

**Nanodisc Reconstitution.** For nanodisc reconstitution, the purified Cx32 protein was mixed with MSP2N2 and lipids, either 1-palmitoyl-2-oleoyl-glycero-3-phosphocholine (POPC) or liver polar lipids (LPL), diluted in buffer C to 200 µl, in a molar ratio of 6:2:200 (Cx32:MSP2N2:lipid) and incubated at room temperature at constant rotation for 1 h. The Bio-Beads SM2 Resin (BioRad, 50 mg) was added to the mixture, placed at 4 °C with constant rotation overnight. The protein was separated from the BioBeads into a fresh Eppendorf tube, centrifuged for 5 min at 16 000 rpm and 4 °C using a cooling benchtop centrifuge (Eppendorf). The reconstituted protein was further purified by HPLC using a Superose 6 Increase 10/300 GL column pre-equilibrated with buffer A. The size exclusion chromatography (SEC) fractions corresponding to Cx32 in nanodisc were pooled together and concentrated to ~2 mg/ml (samples: Cx32 POPC nanodisc, Cx32 LPL nanodisc, W3S POPC nanodisc) or ~1.3 mg/ml (sample: Cx32 noCHS POPC nanodisc) for cryo-EM sample preparation.

**Cryo-EM Sample Preparation and Data Collection.** For cryo-EM sample preparation, 3.5 µl aliquots of concentrated protein reconstituted in nanodiscs were applied to glow discharged Quantifoil Cu R1.2/1.3 200-mesh grid. The grids were blotted for 3 s and plunge-frozen in liquid ethane using Vitrobot Mark IV (Thermo Fisher).

The cryo-EM datasets were collected using a Titan Krios electron microscope (Thermo Fisher), equipped with a K3 direct electron detector camera (Gatan) and a GIF-Quantum energy filter, with a slit width of 20 eV. The defocus range was set from -0.5 µm to -2.5 µm. The data was collected as movies dose-fractionated into 40 frames using EPU 2.0. The exposure time for each micrograph was 0.9 s, with a total dose of 61.6 $e^-/Å^2$ for Cx32 POPC nanodisc dataset, 61.2 $e^-/Å^2$ and 50 $e^-/Å^2$ for Cx32 LPL nanodisc datasets, 55 $e^-/Å^2$ for Cx32 without CHS POPC nanodisc dataset, and 55 $e^-/Å^2$ for W3S POPC nanodisc dataset.

**Cryo-EM Data Processing.** Optics groups were assigned to individual movies using a script provided by Dr. Pavel Afanasyev [2]. The movies were motion-corrected using MotionCor2 [3]. The Cx32 POPC nanodisc and Cx32 LPL datasets were CTF-corrected using Gctf [4], and Cx32 no CHS POPC nanodisc, and W3S POPC nanodisc datasets using CTFFind4 [5]. The two Cx32 LPL nanodisc datasets were merged immediately after CTF refinement. Approximately 1000 particles were picked per dataset, extracted, and subjected to one round of 2D classification in Relion 4.0.1 [6]. The best classes, showing features of GJC and HC, were selected for reference-based autopicking in Relion 4.0.1. The autopicked particles were subjected for several rounds of 2D classification, until clear GJC features were observed. Best particles were selected for 3D classification, using Cx32 GJC in detergent as a reference (low pass filtered to 40 Å) [1], in D6 symmetry. The best class, showing clear GJC features, was then subjected to 3D auto-refinement with imposed symmetry followed by CTF refinement and particle polishing and final refinement. Autorefinement without imposing any symmetry (C1) was used to assess the influence of symmetry on map quality and features. The local resolution was determined using ResMap [7]. The detailed steps of the processing pipeline for each sample are described in detail in Figures S2-3, S7, S9-10 and Table S1.

Protomer-focused classification was performed by symmetry expanding the particles of the final autorefinement job, using 'relion_particle_symmetry_expand' command, using D6 symmetry. The reference map for a single protomer was generated in ChimeraX [8], based on the corresponding model, using the 'Color Zone' and 'Split Map' commands. The reference was used for mask generation in Relion 4.0.1, with 10 Å low pass filtering, extending the binary map with 5 pixels and adding a 20-pixel soft edge. The particles were then subtracted based on the generated mask, recentred on the mask, and rescaled to box size of 200 x 200 pixels. The particles were used for 3D classification without image alignment and symmetry imposition, using appropriately recentred and rescaled mask and reference, to 4, 6 or 8 classes. The best class number was selected by evaluating if more classes reveal more information in the regions of interest without concomitant drastic decrease in resolution (Figure S7).

**Model Building and Refinement.** The models of Cx32 GJC in POPC nanodisc, Cx32 without CHS GJC in POPC GJC nanodisc, and Cx32 GJC in LPL nanodisc were built based on the single subunit of Cx32 WT HC detergent model (PDB ID: 7ZXN). The models of W3S GJCC in POPC nanodisc and

R22G GJC in POPC nanodisc were built based on the single subunit of W3S and R22G HC detergent models, respectively (PDB ID: 7ZXT and 7ZXO). The models were built in COOT [9] by aligning the subunit template into the density and refining amino acid positions. The cytoplasmic regions (CL: L106-H126 (Cx32 POPC, Cx32 LPL, W3S POPC); CT: A218-C283 (Cx32 POPC, Cx32 LPL, W3S POPC) were not built due to not being resolved in the density map. The cholesterol (lipid2) (chemical ID: CLR) was built into Cx32 POPC nanodisc maps, based on Cx32 HC detergent model (PDB ID: 7ZXN). POPC molecule (chemical ID: POV) was built into Cx32 POPC and LPL nanodisc maps, using first rigid body fit and later real space refine functions in COOT. The side-chain clashes and overlaps were minimized using Chiron [10]. All models were refined using 'phenix.real_space_refine' function and validated using MolProbity [11] in PHENIX [12]. The HOLE analysis of the pore radius was performed in COOT.

**Electrostatic Surface Potential Analysis.** The electrostatic potential was calculated in PyMOL using APBS Tools 2.1, using AMBER99 force field.

**Molecular dynamics system setup**

The coordinates for a Cx32 wt connexon bound to six POPC and six CHOL (Cx32 wt:POPC:CHOL) molecules were extracted from the cryo-EM structure. The protonation states of the protein's ionizable residues and the transmembrane region were predicted using the H++ and PPM web servers, respectively [13,14]. ACE and NME caps were added to residues M105 and I127, respectively, in each Cx32 chain, corresponding to gaps found in the cryo-EM structures.

CHARMM-GUI [15] was employed to embed the protein into a lipid bilayer containing major components of plasma membranes from mammalian cells, i.e., 1-palmitoyl-2-oleoylphosphatidylcholine (POPC), 1-palmitoyl-2-oleoylphosphatidylethanolamine (POPE), palmitoyl sphingomyelin (PSM), 1-palmitoyl-2-oleoylphosphatidylserine (POPS) and cholesterol (CHOL), that are parametrized in the lipid21 force field of Amber 22 (https://opm.phar.umich.edu/biological_membranes/lipid_composition). The resulting bilayer comprised a total of 146 CHOL molecules, 126 POPC molecules, 68 PSM molecules, 68 POPE molecules and 22 POPS molecules. Water molecules were added to form a cuboid box measuring 134.37 Å x 135.51 Å x 102.23 Å, with edges positioned at least 10 Å away from the protein surface and the bilayer stretching along the xy plane.

The system's PDB file, assembled using CHARMM-GUI, was processed by the program charmmlipid2amber.py from AmberTools 22 in order to convert the CHARMM naming convention into AMBER [16]. The PDB files for other systems prepared for MD simulations, which included Cx32 wt with no lipids in the pore (apo Cx32 wt), apo Cx32 W3S, Cx32 W3S with POPC and CHOL in the pore (Cx32 W3S:POPC:CHOL), Cx32 wt with only POPC in the pore (Cx32 wt:POPC), Cx32 wt with only CHOL in the pore (Cx32 wt:CHOL), Cx32 W3S with only POPC in the pore (Cx32 W3S:POPC) and Cx32 W3S with only CHOL in the pore (Cx32 wt:CHOL), were generated from the processed PDB of Cx32 wt:POPC:CHOL, either the initial structure or an equilibrated frame (see below). In all cases, "CONECT" records for S-S bonds in the proteins were obtained using *pdb4amber* of Amber 22 [16], which ensured the correct formation of such bonds before further processing for MD simulations.

The program *tleap* was employed to obtain the topologies and the coordinate files for the studied systems and to add the counterions ($Na^+$) necessary to neutralize the simulation boxes. The force fields ff19SB and lipid21 were chosen to parametrize the protein and the lipids, including those in the bilayer or within the protein pore. The water molecules were treated with the OPC model [16]. Hydrogen mass repartitioning was carried out to increase the time step from 2 to 4 fs during the production runs [17].

### Molecular dynamics simulations

Prior to the production runs, all systems were subjected to energy minimization (EM) and equilibration steps. A total of 3,000 EM cycles were performed for each system using the program *pmemd.MPI* [16]. The minimized systems were subjected to heating in two subsequent simulations. During the first phase, the temperature was linearly increased from 10 to 100 K during 500 ps. Then, during the second phase, the temperature was linearly increased from 100 to 303 K during 600 ps. In both cases, the heavy atoms of the protein and all lipids were restrained to their initial positions using a harmonic potential defined by a spring constant of 10 kcal·$Å^{-2}$·$mol^{-1}$. The cell volume was kept constant and the temperature was controlled using the Langevin thermostat, with a collision frequency of 1.0 $ps^{-1}$.

Following the heating phase, the systems underwent four NPT equilibration steps, during which position restraints were applied exclusively to the heavy atoms of the protein and the lipids within the pore (if present). The restraint constant was reduced from 8 to 2 kcal·$Å^{-2}$·$mol^{-1}$ in 2 kcal·$Å^{-2}$·$mol^{-1}$ strides over the four 8 ns MD simulations. During these equilibration steps, the systems were simulated at constant temperature and pressure of 303 K and 1 bar, which was achieved by using the Langevin thermostat and the Monte Carlo barostat, respectively. It is worth noting that the initial structures to generate the topology and coordinate files of systems involving the W3S mutation were derived from the equilibrated structures of the wild-type systems. Finally, all systems were subjected to 1 µs production runs in the NPT ensemble, in identical conditions to those used during the equilibration except for the position restraints, which were eliminated. All the steps, from heating to production runs, were carried out in duplicate with *pmemd.cuda* [18].

### Trajectory analysis

Structural analyses along the trajectories were carried out with different commands of the program *cpptraj* of Amber22 [16]. The *secstruct* and *rmsf* commands were employed to monitor the secondary structure and atomic fluctuations of residues within the N-terminal region of Cx32 along the MD trajectories, respectively. Moreover, the *rms and angle* commands were used to calculate root-mean-square deviations for the bound lipids and to monitor the angle formed between the NTH and TM1 during the MD simulations, respectively [16].

Clustering analysis was performed with the *cluster* command of *cpptraj* to determine the representative structure of Cx32 and the bound lipids during the MD simulations. The hierarchical agglomerative method was chosen for clustering and the RMSD the heavy atoms of the bound lipids and of residues belonging to the lipid binding site were used as the metric for clustering [16].

### Supplementary Figures

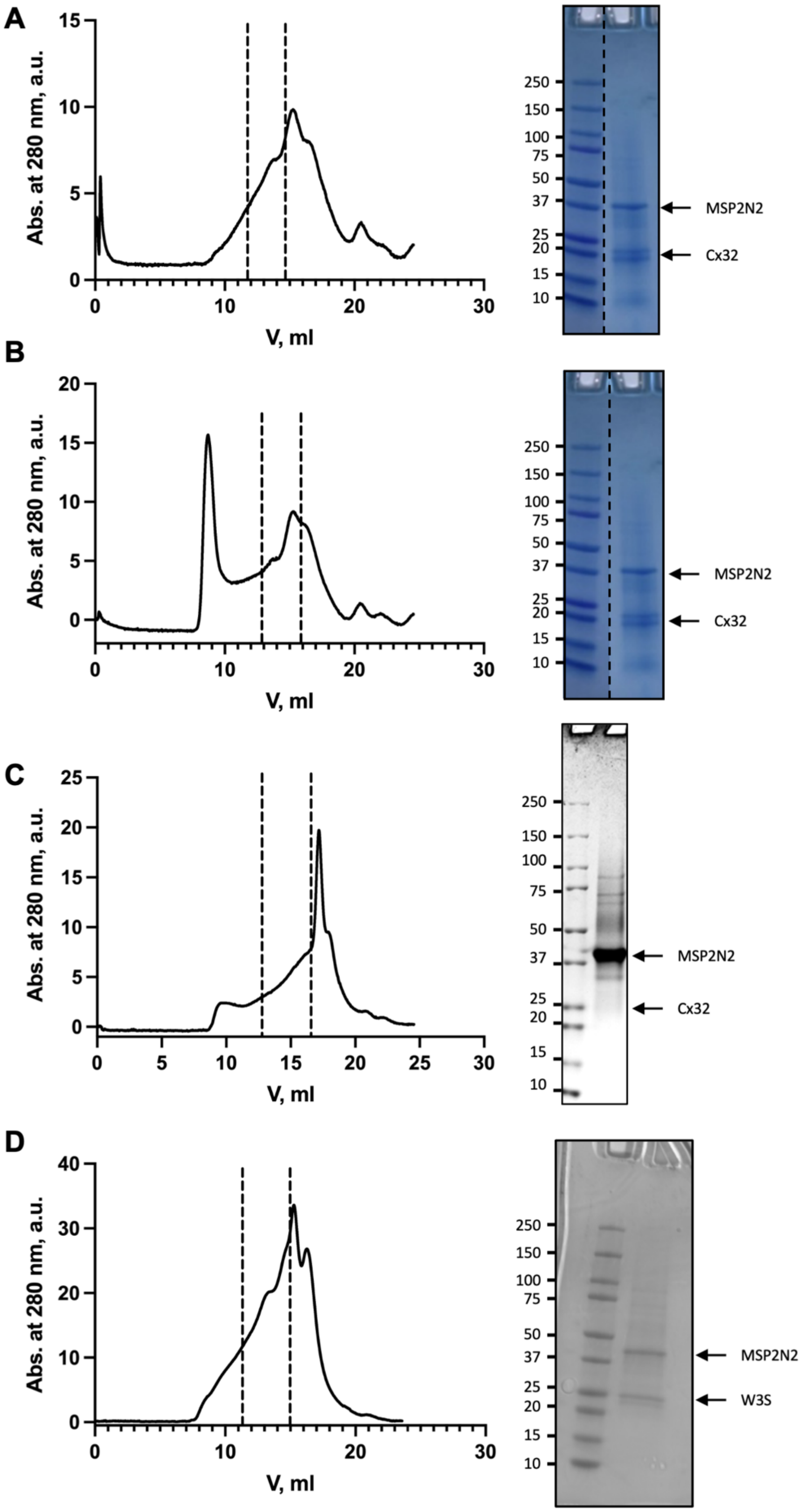

**Figure S1. Size exclusion chromatograms and SDS PAGE gels of nanodisc reconstitution** of **(A)** Cx32 in POPC, **(B)** Cx32 in LPL, **(C)** W3S in POPC, and **(D)** Cx32, purified in the absence of CHS, in POPC nanodisc, using MSP2N2 as the nanodisc scaffold protein.

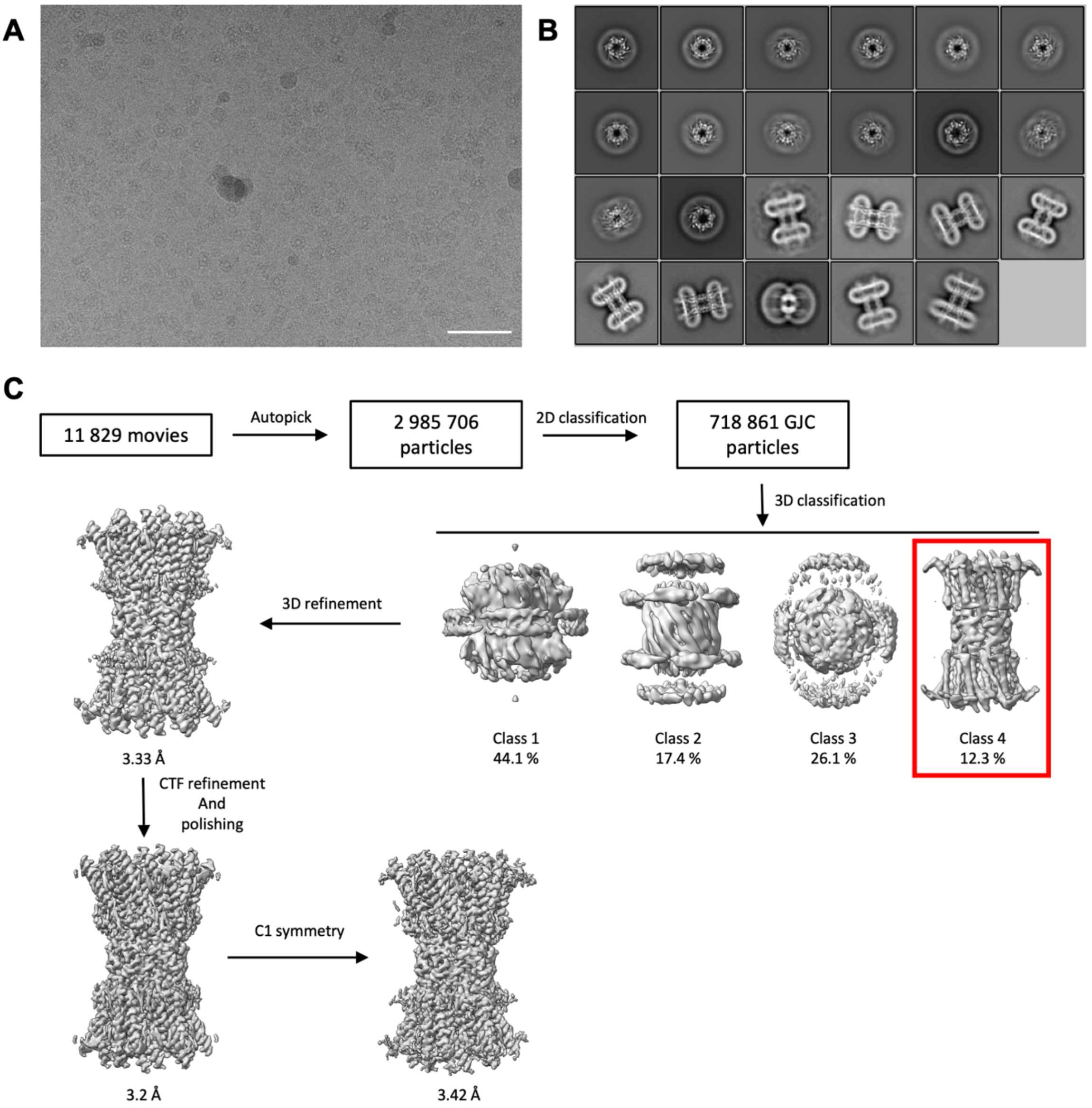


**Figure S2. Cryo-EM structure determination of Cx32 GJC in POPC nanodisc. (A)** Cryo-EM micrograph of Cx32, reconstituted in POPC-containing nanodisc. Scale bar = 50 nm. **(B)** 2D classes of Cx32 GJC particles, used for 3D classification. **(C)** Cryo-EM data processing pipeline for Cx32 GJC in POPC nanodisc.

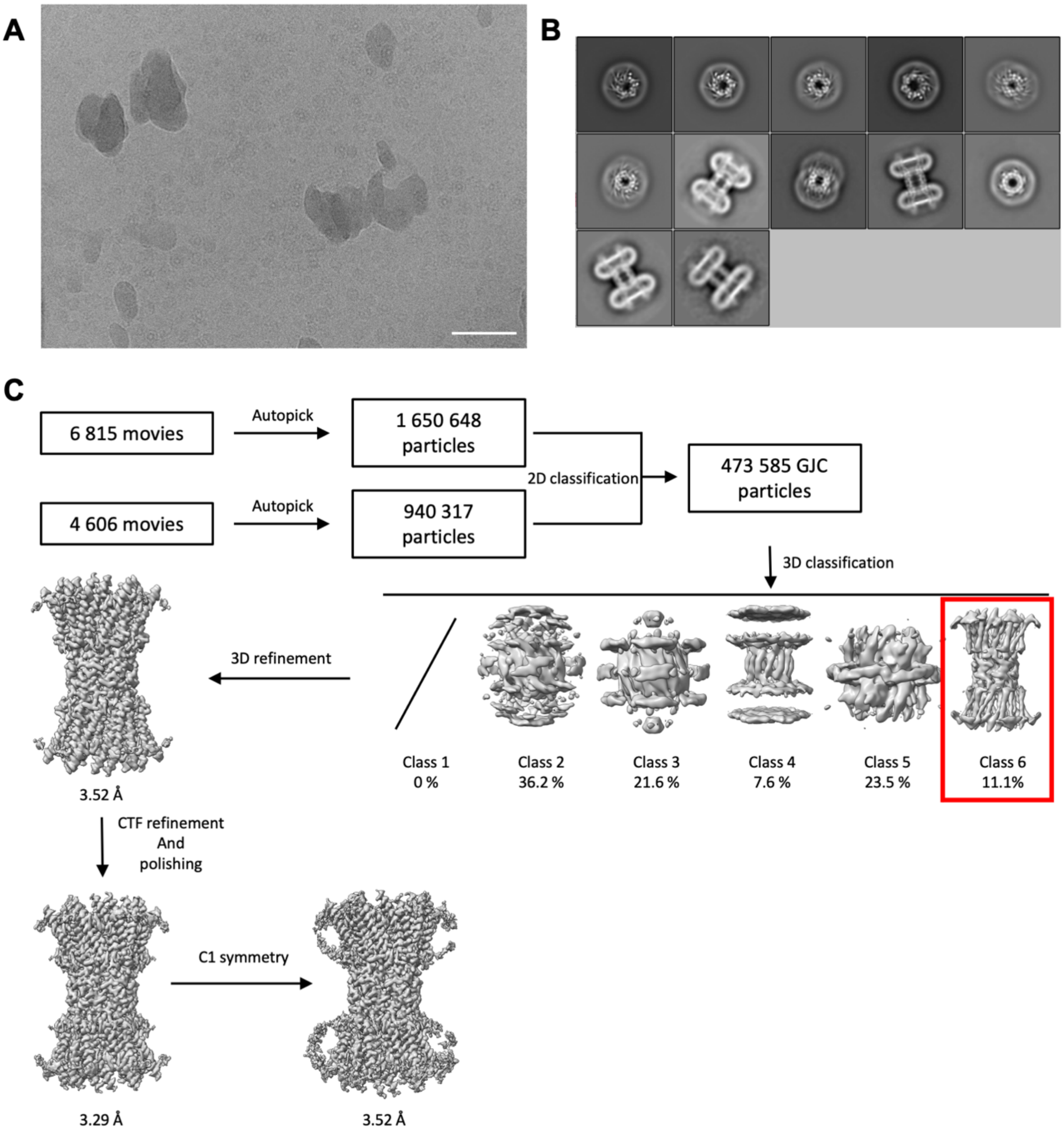


**Figure S3. Cryo-EM structure determination of Cx32 GJC in LPL nanodisc. (A)** Cryo-EM micrograph of Cx32, reconstituted in LPL-containing nanodisc. Scale bar = 50 nm. **(B)** 2D classes of Cx32 GJC particles, used for 3D classification. **(C)** Cryo-EM data processing pipeline for Cx32 GJC in LPL nanodisc.

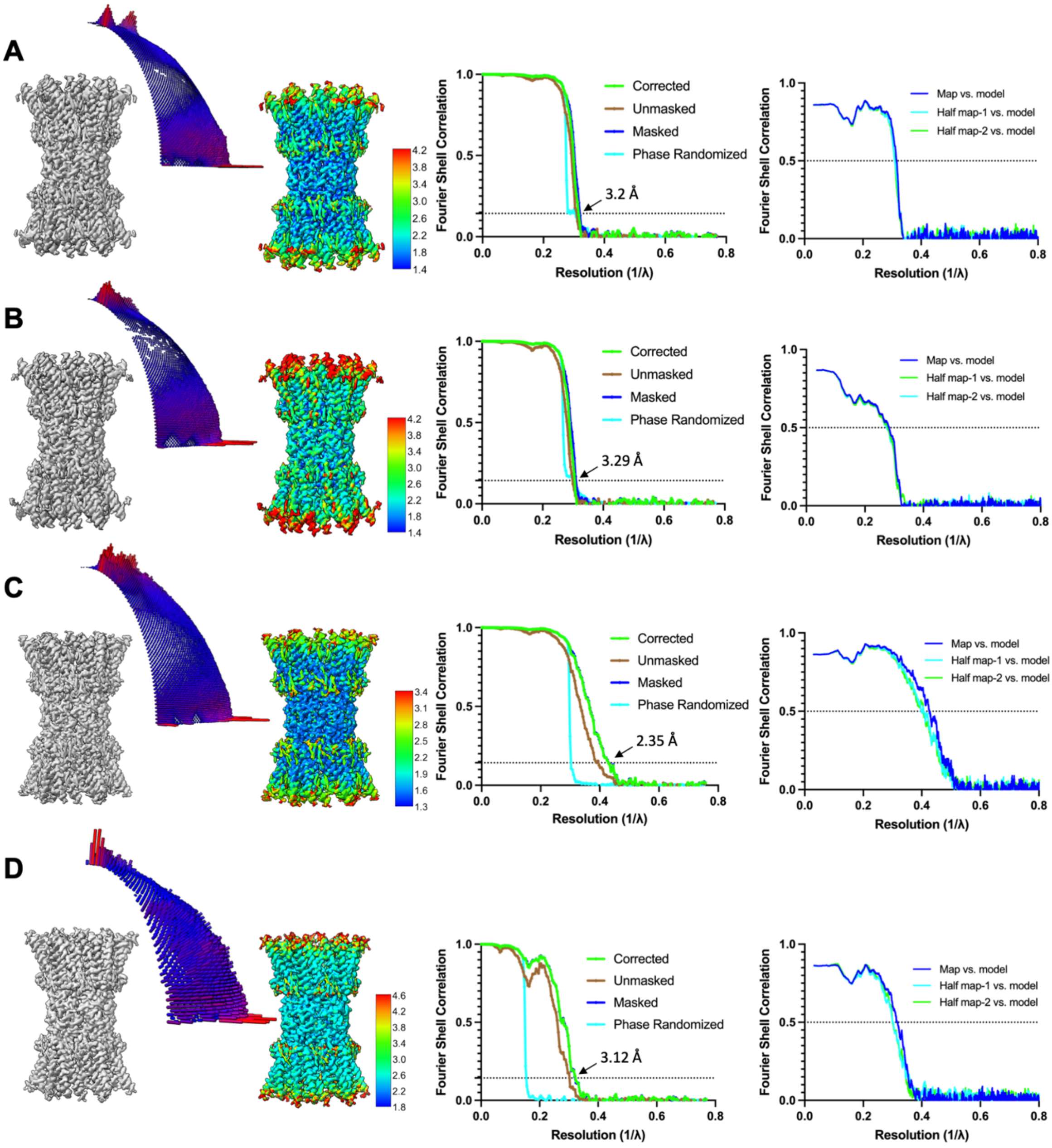


**Figure S4. Angular distribution, local resolution, and Fourier Shell Correlation (FSC) curves.** **(A)** Cx32 GJC in POPC nanodisc. **(B)** Cx32 GJC in LPL nanodisc. **(C)** W3S GJC in POPC nanodisc. **(D)** Cx32 GJC, without CHS, in POPC nanodisc.

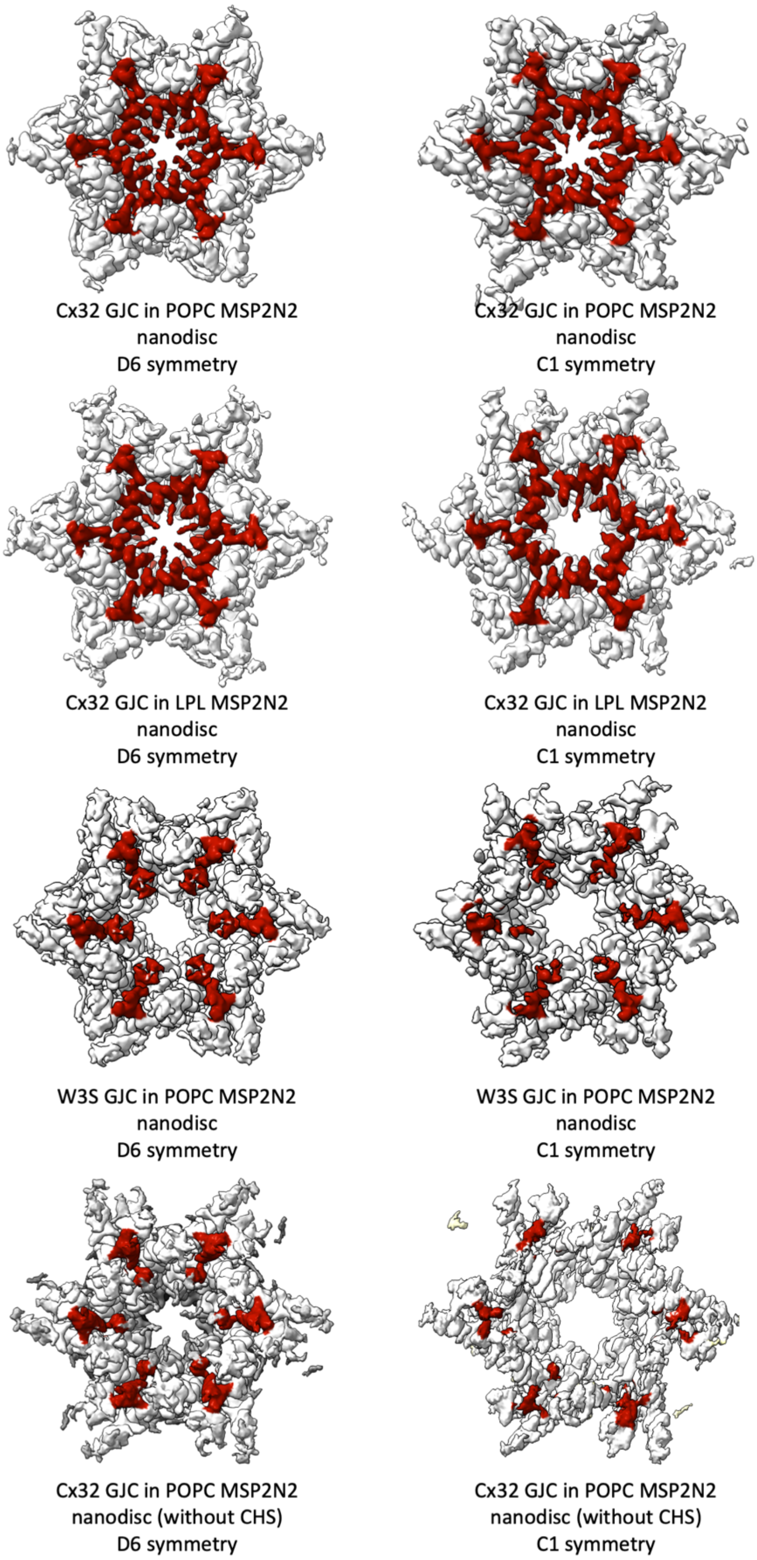


**Figure S5. Top view comparison of Cx32 GJC nanodisc maps in D6 and C1 symmetry.** The densities corresponding to the N-terminus are represented in red.

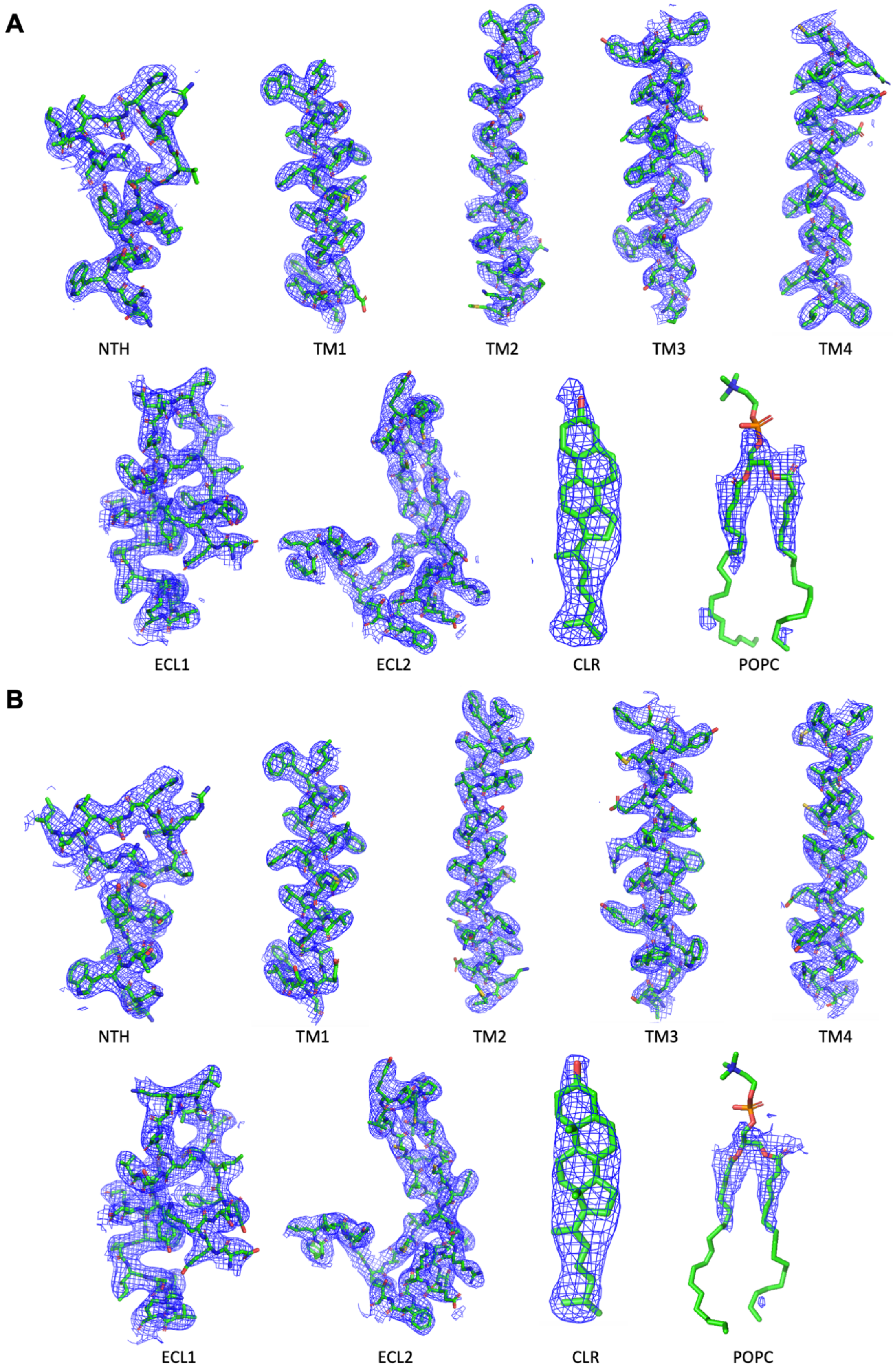


**Figure S6. Cryo-EM density map features of Cx32 GJC in (A) POPC-containing and (B) LPL-containing nanodiscs.**

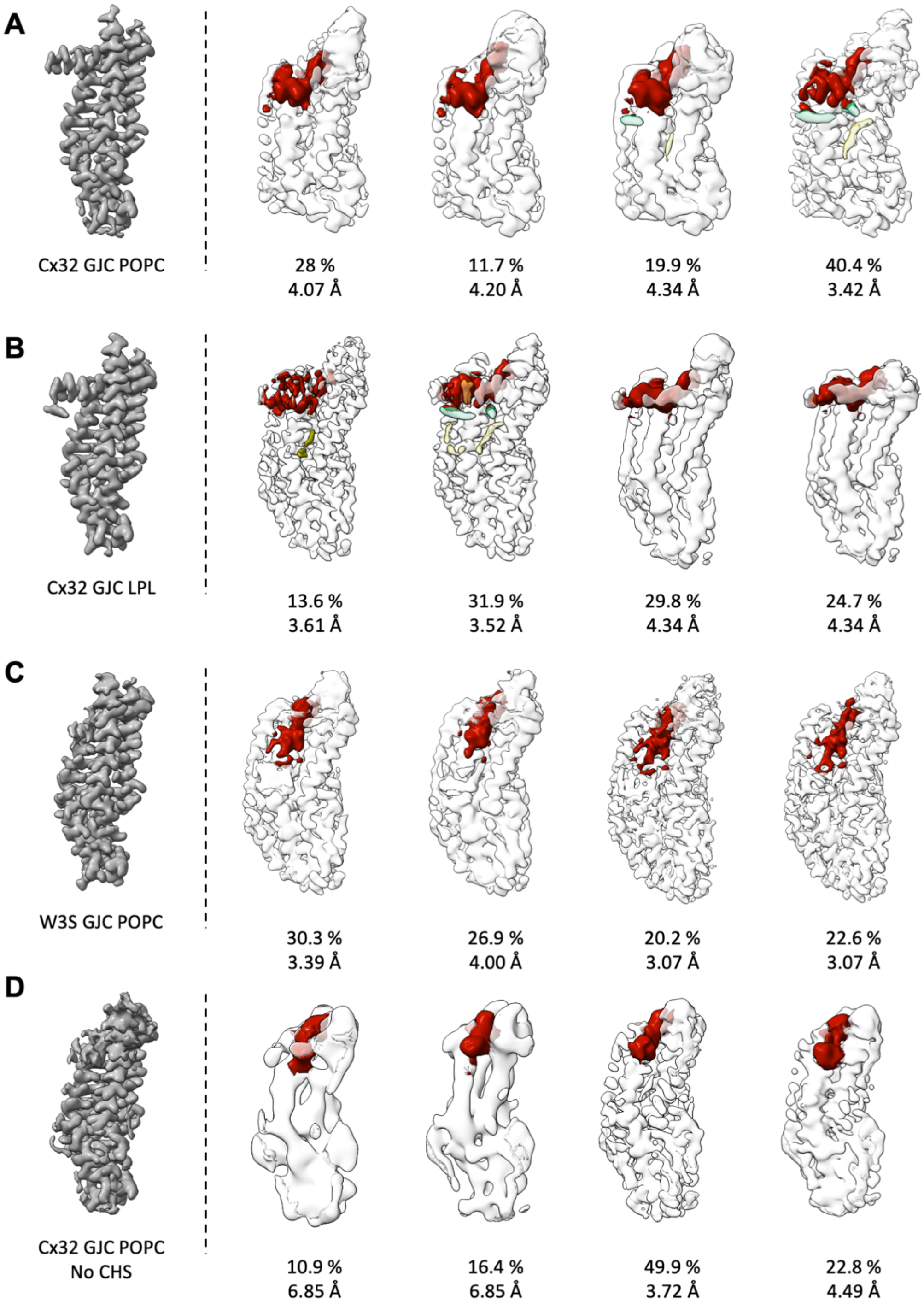


**Figure S7. Protomer-focused classification. (A)** Cx32 GJC in POPC nanodisc. **(B)** Cx32 GJC in LPL nanodisc. **(C)** W3S GJC in nanodisc. **(D)** Cx32 GJC, without CHS, in nanodisc. Red coloured density corresponds to the NTH.

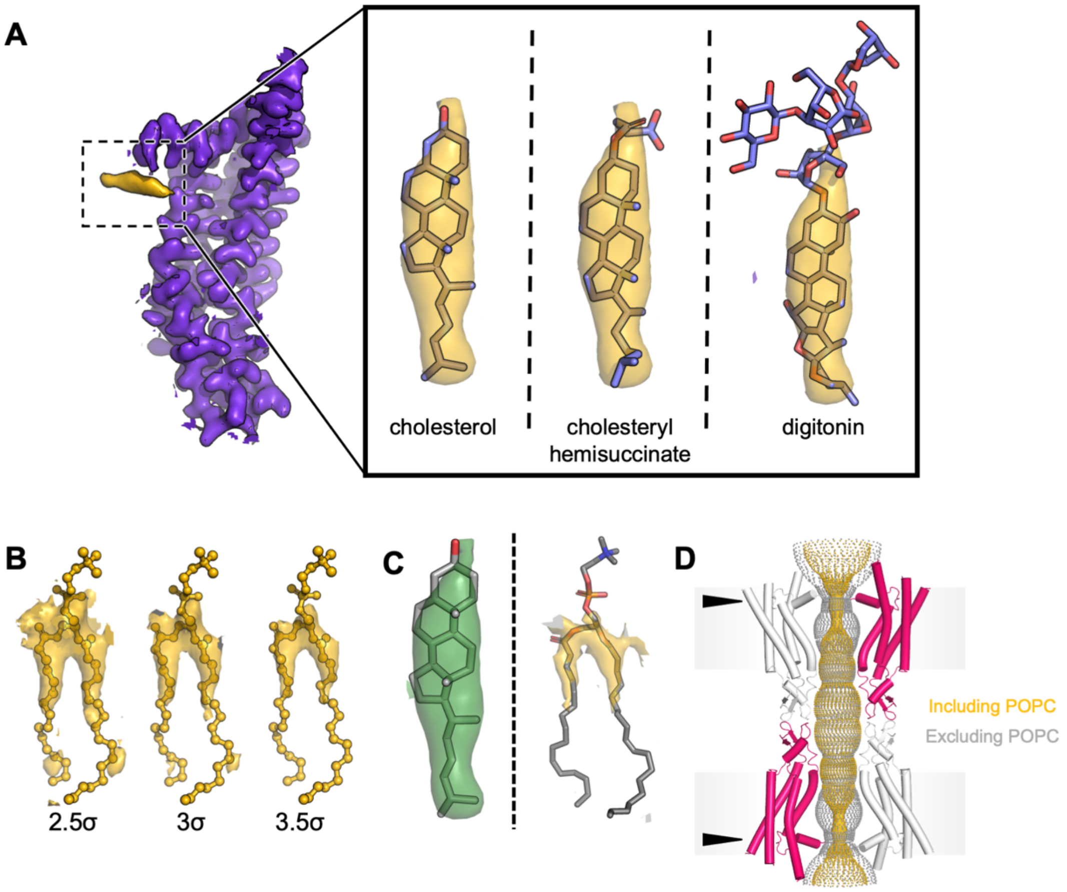


**Figure S8. Effect of lipid-2 on Cx32 GJC NTH. (A)** Lipid-2 density can accommodate cholesterol, or cholesteryl hemisuccinate (CHS), and digitonin, used during protein purification. **(B)** Cx32 GJC reconstituted in LPL-containing nanodisc has the similar densities as Cx32 GJC, reconstituted in POPC containing nanodisc. **(C)** HOLE analysis of the pore conduction pathway of Cx32 GJC in LPL-containing nanodiscs. The pathway colored in yellow includes POPC in the calculation, whereas the grey excludes it. The arrows represent the points of pore constriction due to NTH rearrangement.

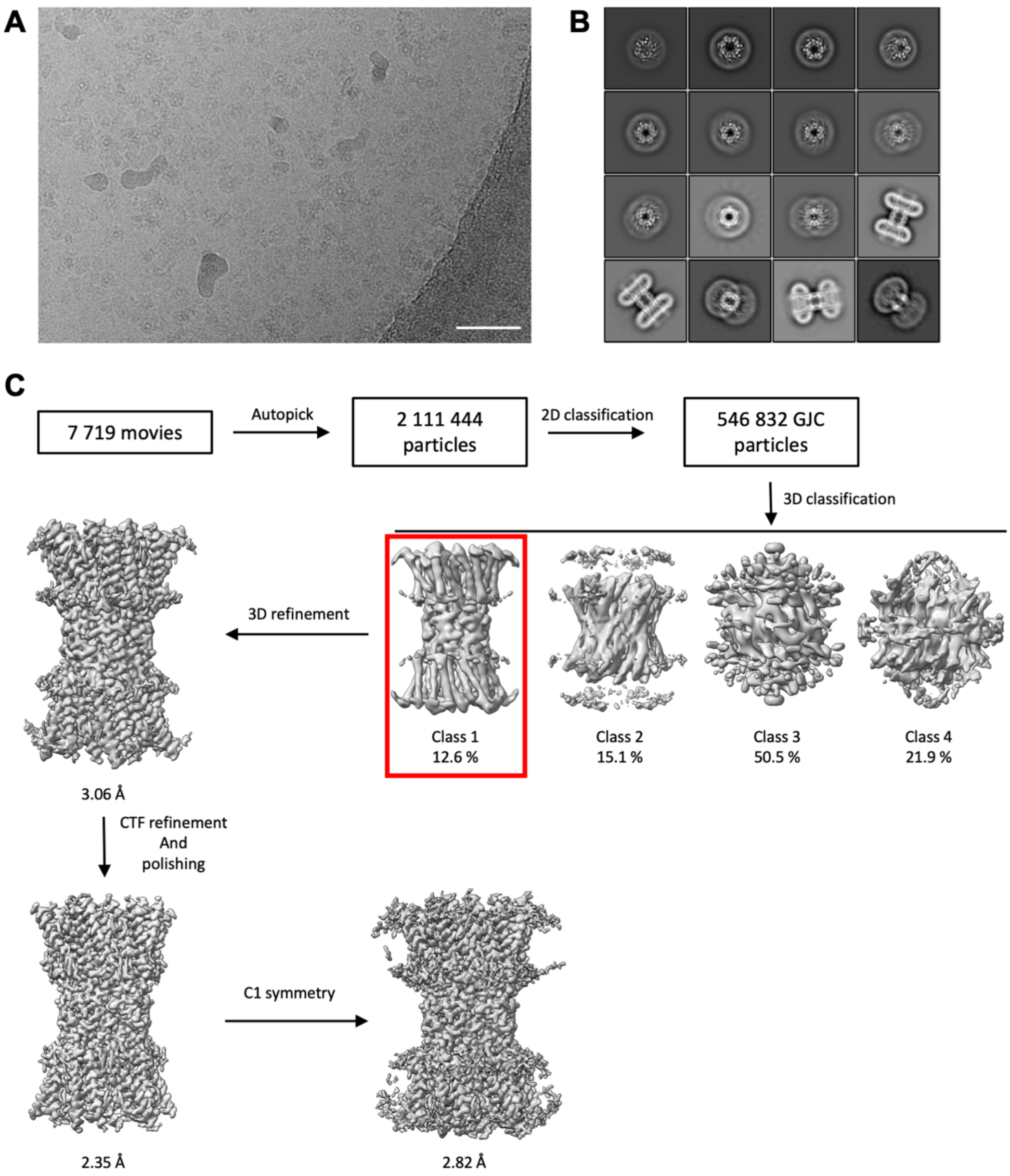


**Figure S9. Cryo-EM structure determination of W3S GJC in nanodisc. (A)** Cryo-EM micrograph of W3S, reconstituted in nanodisc. Scale bar = 50 nm. **(B)** 2D classes of Cx32 GJC particles, used for 3D classification. **(C)** Cryo-EM data processing pipeline for W3S GJC in nanodisc.

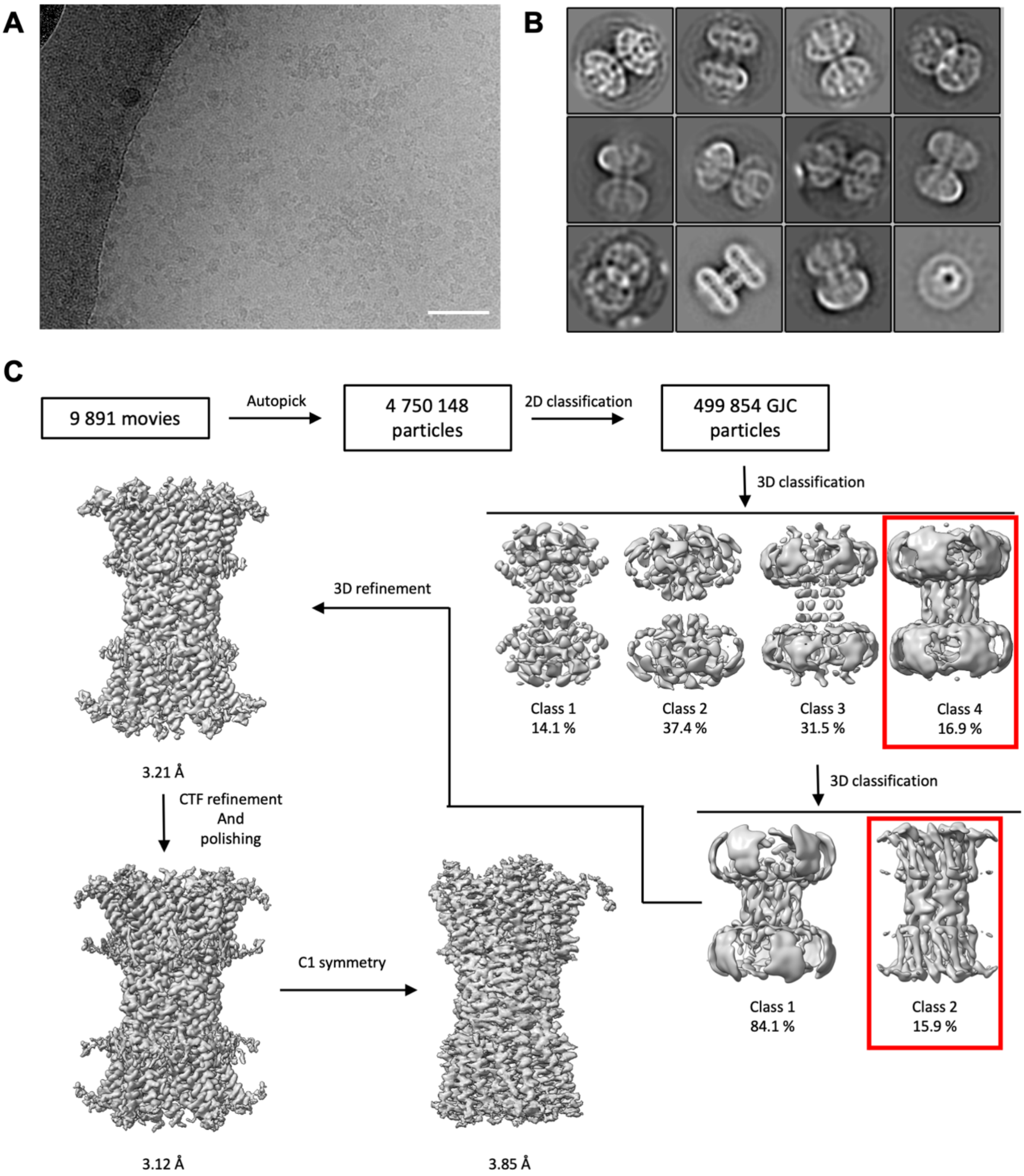


**Figure S10. Cryo-EM structure determination of Cx32 GJC, purified in the absence of CHS, in nanodisc. (A)** Cryo-EM micrograph of Cx32, purified without CHS, in nanodisc. Scale bar = 50 nm. **(B)** 2D classes of Cx32 GJC particles, used for 3D classification. **(C)** Cryo-EM data processing pipeline for Cx32 GJC, purified without CHS, in nanodisc.

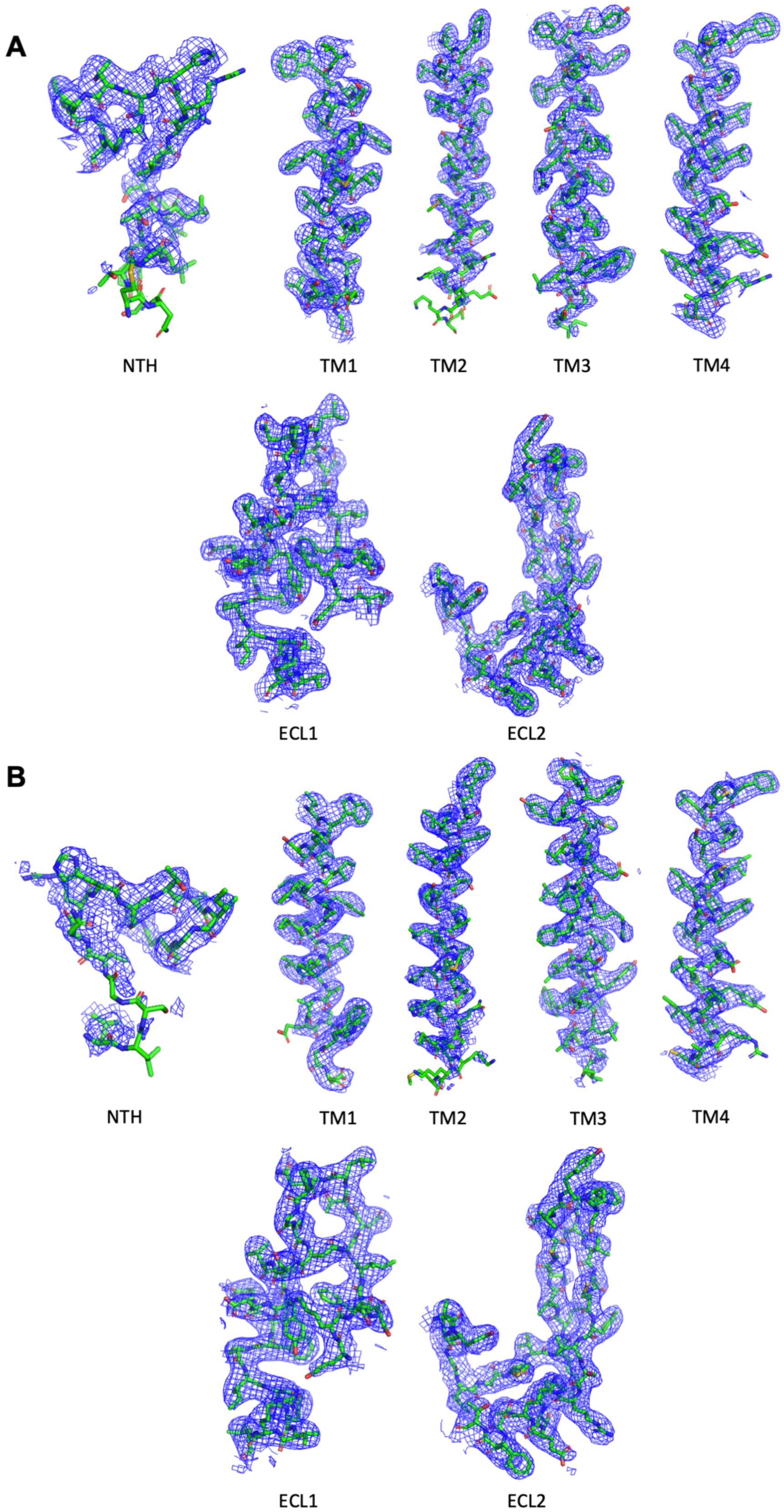


**Figure S11. Cryo-EM density map features of (A) W3S GJC and (B) Cx32 GJC without CHS, in nanodisc.**

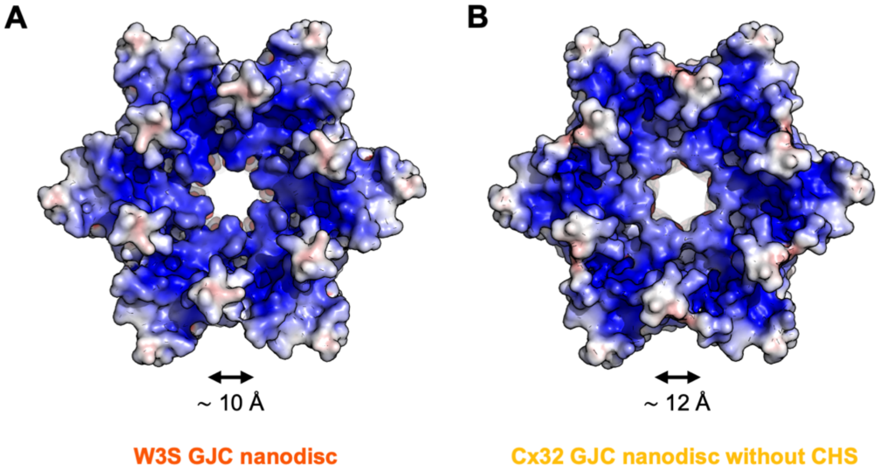


**Figure S12. Electrostatic surface potential of (A) W3S GJC and (B) Cx32 GJC, without CHS, in nanodisc.**

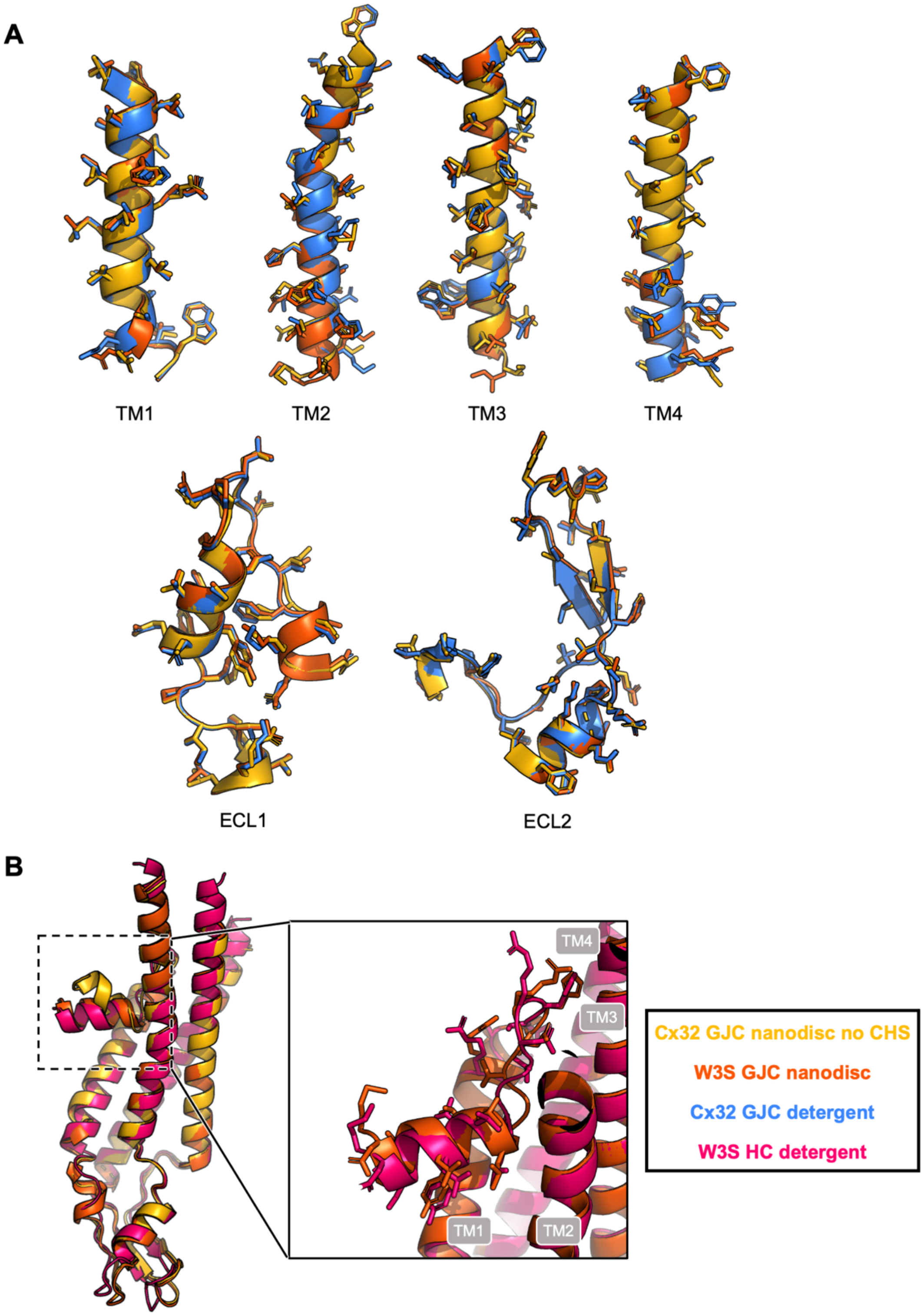


**Figure S13. Comparison of W3S and Cx32, without CHS, GJC structures in detergent and nanodisc. (A)** Comparison of TMH and ECL of W3S and Cx32, without CHS, GJC in nanodiscs compared to Cx32 GJC structure in detergent. **(B)** Comparison of W3S GJC in nanodisc to W3S HC in detergent.

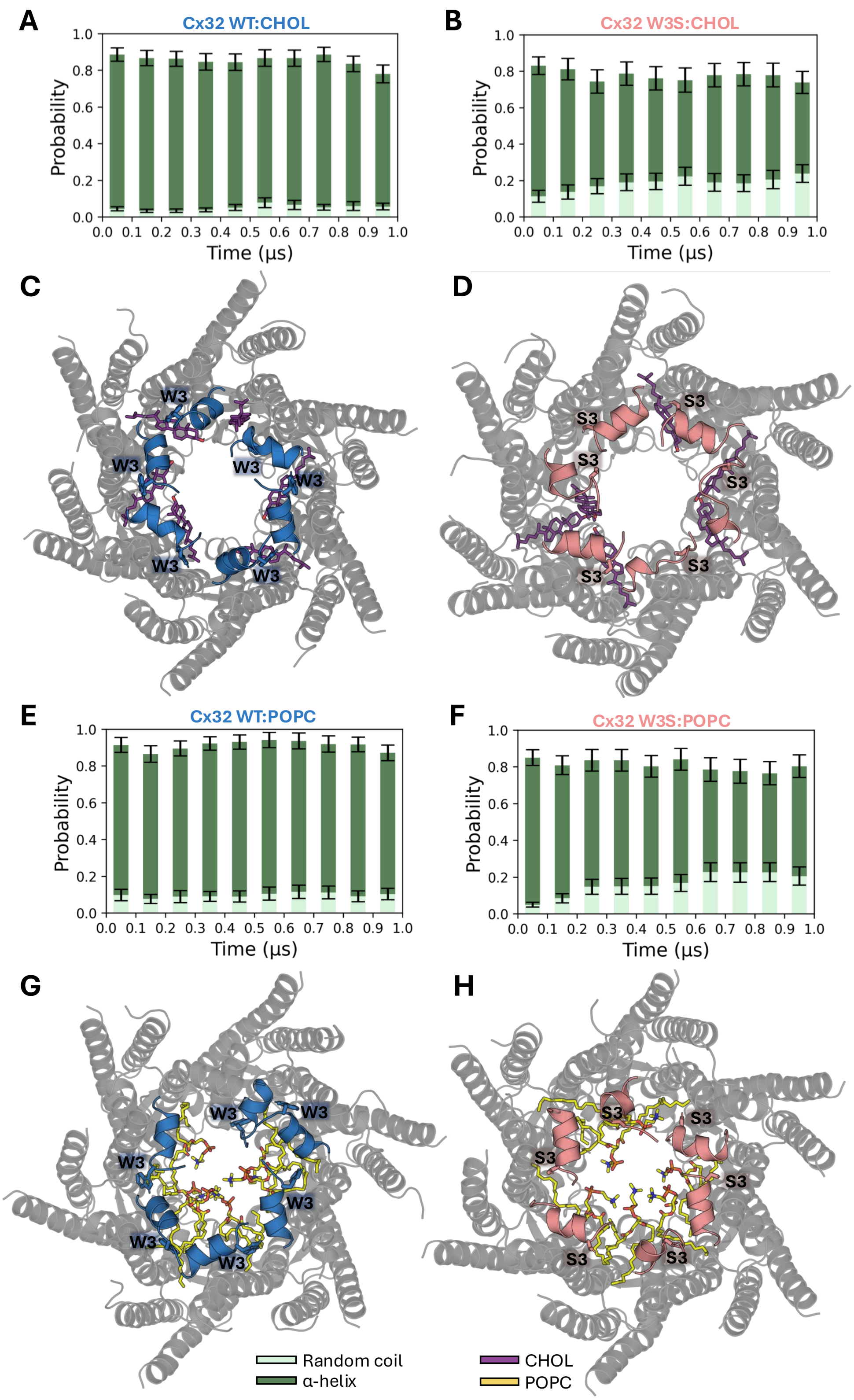


**Figure S14. Time evolution of the secondary structure of residues 3-10 in Cx32 bound to either CHOL or POPC during 1 µs MD simulations. (A-B), (E-F)** represent the mean probabilities of residues 3-10 (which form the N-terminal α-helices in Cx32 wt) adopting an α-helix or random coil conformation for the indicated systems throughout the simulation time. For each system, the plotted mean probabilities were calculated by averaging the per-residue probabilities for residues within the 3-

10 interval in every protein chain and over the two replicate MD simulations in 0.1 μs intervals (i.e., 0-0.1, ..., 0.9-1 μs). Other types of secondary structure are omitted. **(C-D)**, **(G-H)** show a structural representation of the last frame (*t*=1 μs) in one of the two replicate trajectories for each system. The N-terminal residues 1-10 are colored in blue and salmon in each panel.

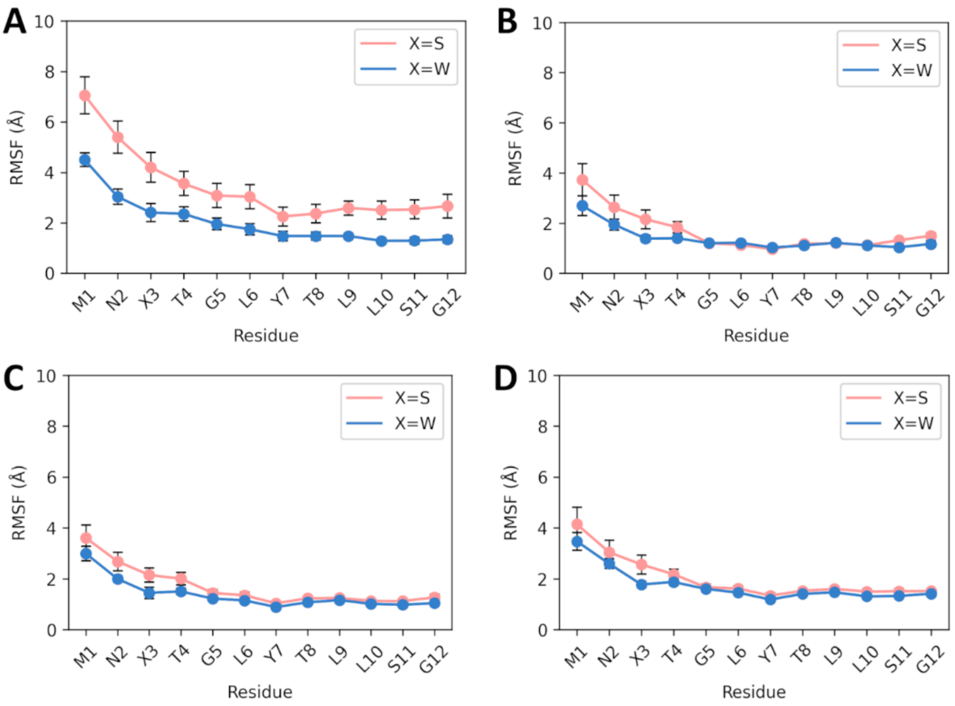


**Figure S15. Root-mean-square fluctuation (RMSF) values for the N-terminal residues 1-12 of Cx32 wt and Cx32 W3S in the presence and absence of lipids in the pore. (A)** Cx32 wt and Cx32 W3S, (**B)** Cx32 wt:POPC:CHOL and Cx32 W3S:POPC:CHOL, (**C)** Cx32 wt:CHOL and Cx32 W3S:CHOL, and (**D)** Cx32 wt:POPC and Cx32 W3S:POPC. Average RMSF values were calculated for residues 1-12 in the six chains of each protein and during two replicate 1 μs MD simulations for each system. The first 100 ns of each MD trajectory were excluded from the RMSF analysis. All fluctuations were calculated with respect to the energy-minimized structure of each system and considering the heavy atoms of the selected residues.

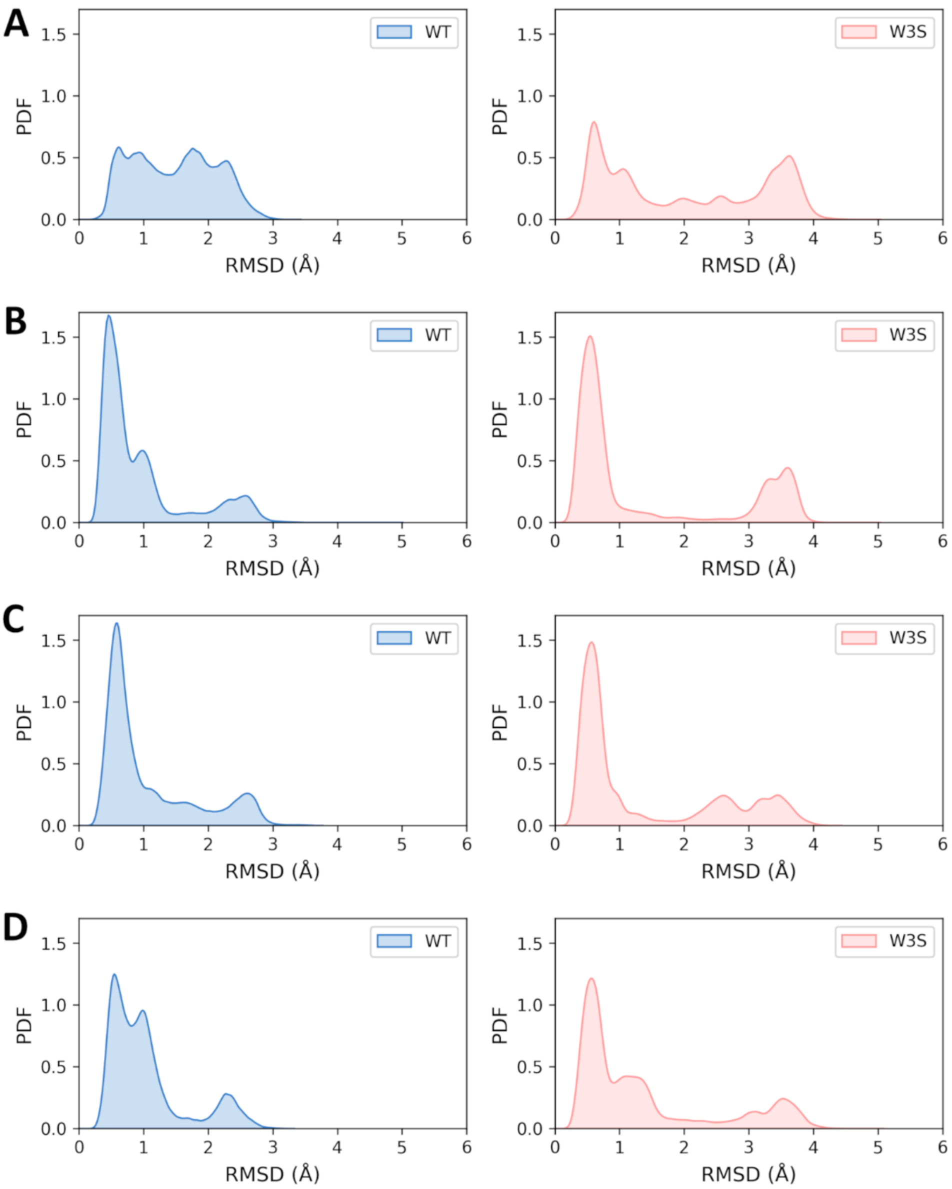


**Figure S16. Distributions of root-mean-square deviation (RMSD) values for the N-terminal residues 1-10 of Cx32 wt and Cx32 W3S in the presence and absence of lipids in the pore. (A)** Cx32 wt and Cx32 W3S, **(B)** Cx32 wt:POPC:CHOL and Cx32 W3S:POPC:CHOL, **(C)** Cx32 wt:CHOL and Cx32 W3S:CHOL, and **(D)** Cx32 wt:POPC and Cx32 W3S:POPC. Average RMSD values were calculated for the backbone of residues 1-10 in the six chains of each protein and during two replicate 1 μs MD simulations for each system. PDF on the *y* axis stands for probability density function. RMSD values for each N-terminal segment (residues 1-10) were calculated relative to the position of the corresponding segment in the energy-minimized structure of each system, with trajectories fitted onto the initial backbone positions of these residues and after discarding the first 100 ns. Thus, these RMSD calculations focus on the internal motion of the N-terminal residues without reflecting their movement relative to the rest of the protein.

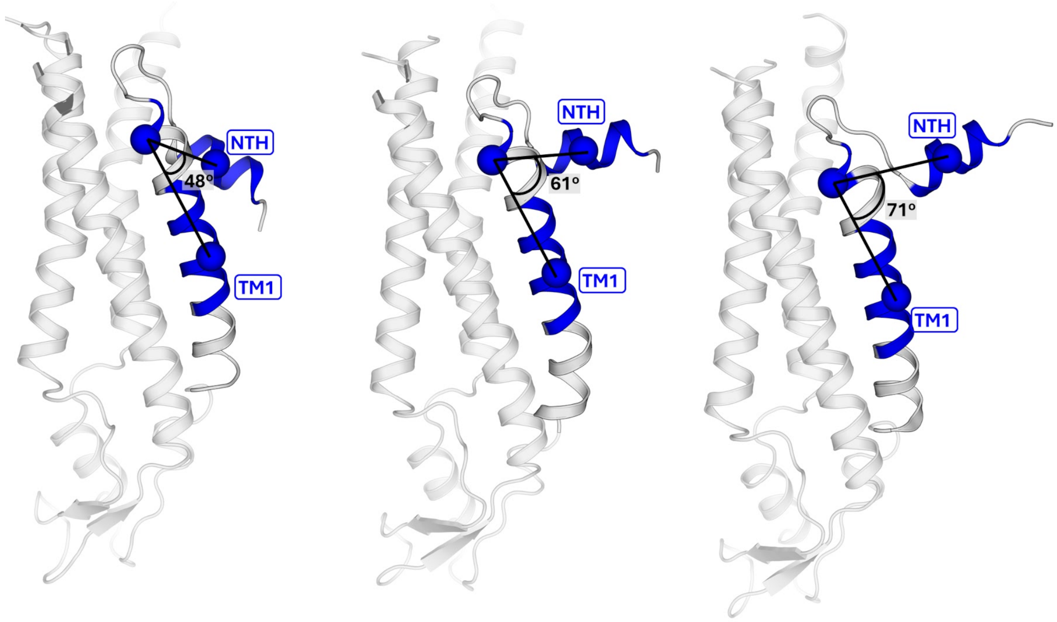


**Figure S17.** Definition of the angle used to monitor NTH motion relative to TM1. The centers of mass (blue spheres) of three groups of Cα atoms, corresponding to residues 3–10, 19–21, and 26–35 in each chain (represented by blue segments), were used to define the NTH-TM1 angle. In the cryo-EM structure of Cx32 in nanodiscs, this angle is approximately 61° (center structure). During MD simulations, both smaller and larger angles are sampled (see left and right structures).

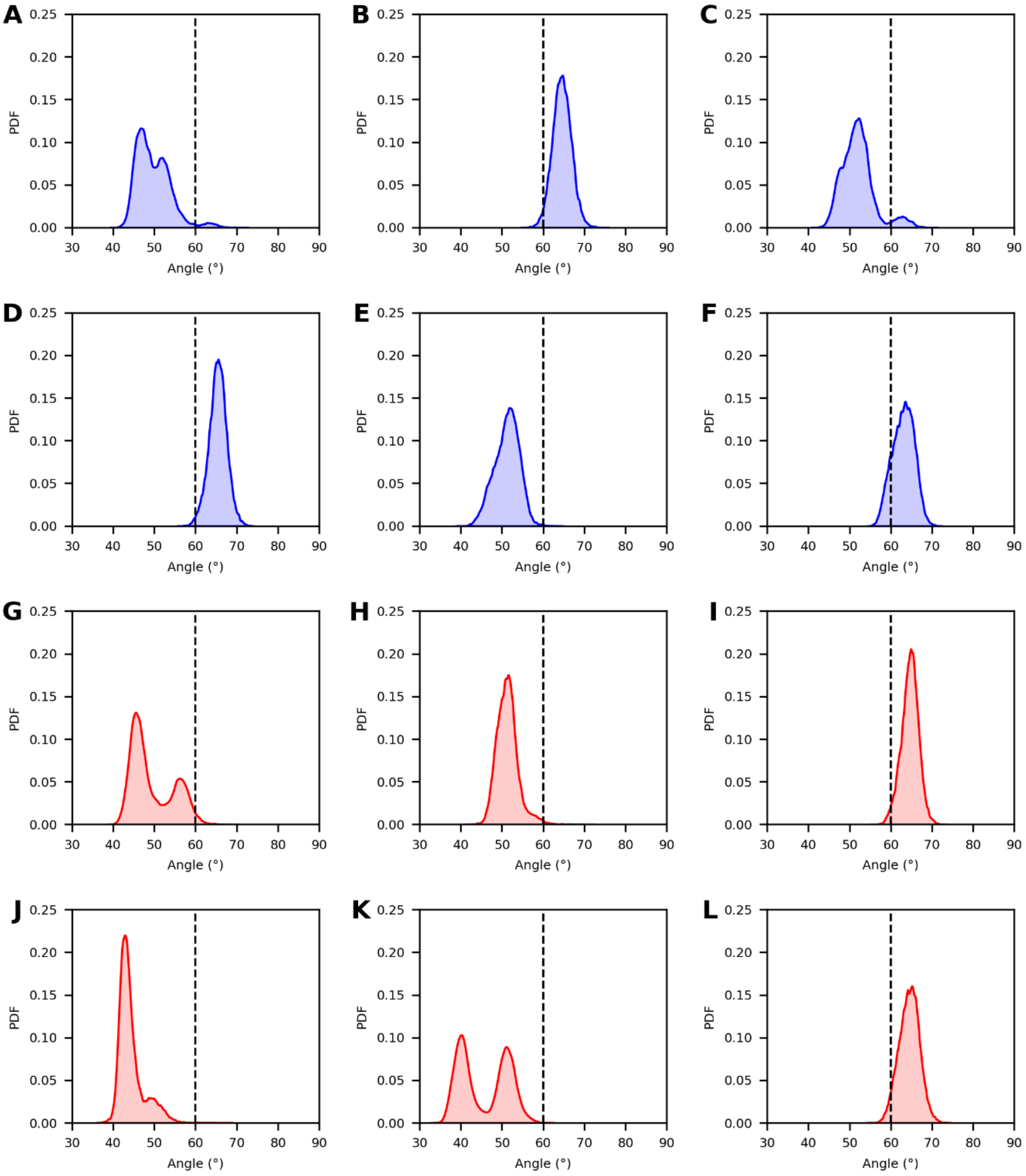


**Figure S18.** Distributions of NTH-TM1 angles in Cx32 wt. **(A–F)** Angle distributions for each chain of the Cx32 hexamer during the first replicate MD simulation (blue). **(G–L)** Angle distributions for each chain during the second replicate MD simulation (red). Initial and final panels (A, F and G, L), and adjacent panels represent neighbouring chains in the structure. The vertical dashed line indicates the value of the angle in the cryo-EM structure of Cx32 in nanodiscs.

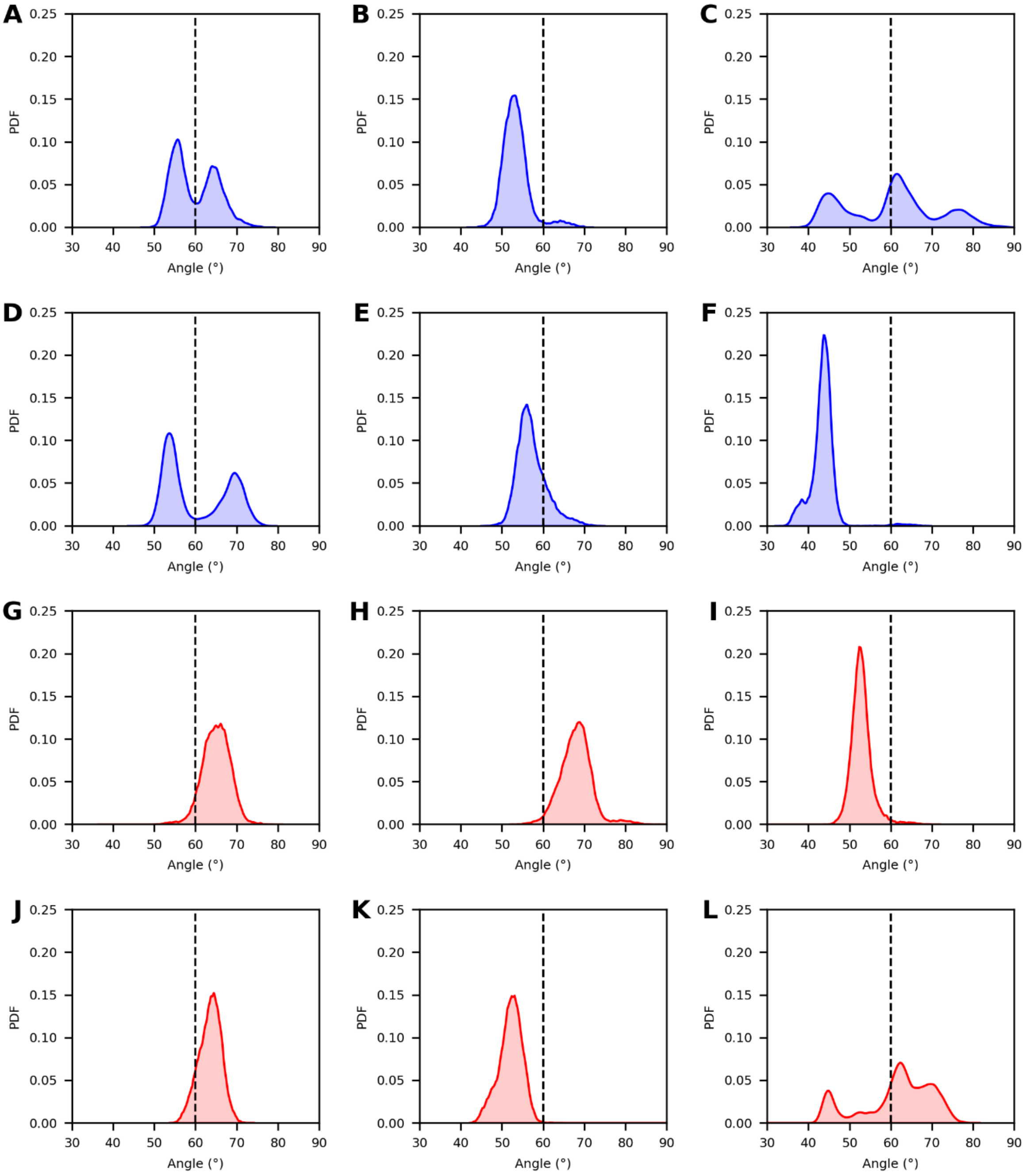


**Figure S19.** Distributions of NTH-TM1 angles in Cx32 W3S. **(A–F)** Angle distributions for each chain of the Cx32 hexamer during the first replicate MD simulation (blue). **(G–L)** Angle distributions for each chain during the second replicate MD simulation (red). Initial and final panels (A, F and G, L), and adjacent panels represent neighbouring chains in the structure. The vertical dashed line indicates the value of the angle in the cryo-EM structure of Cx32 in nanodiscs.

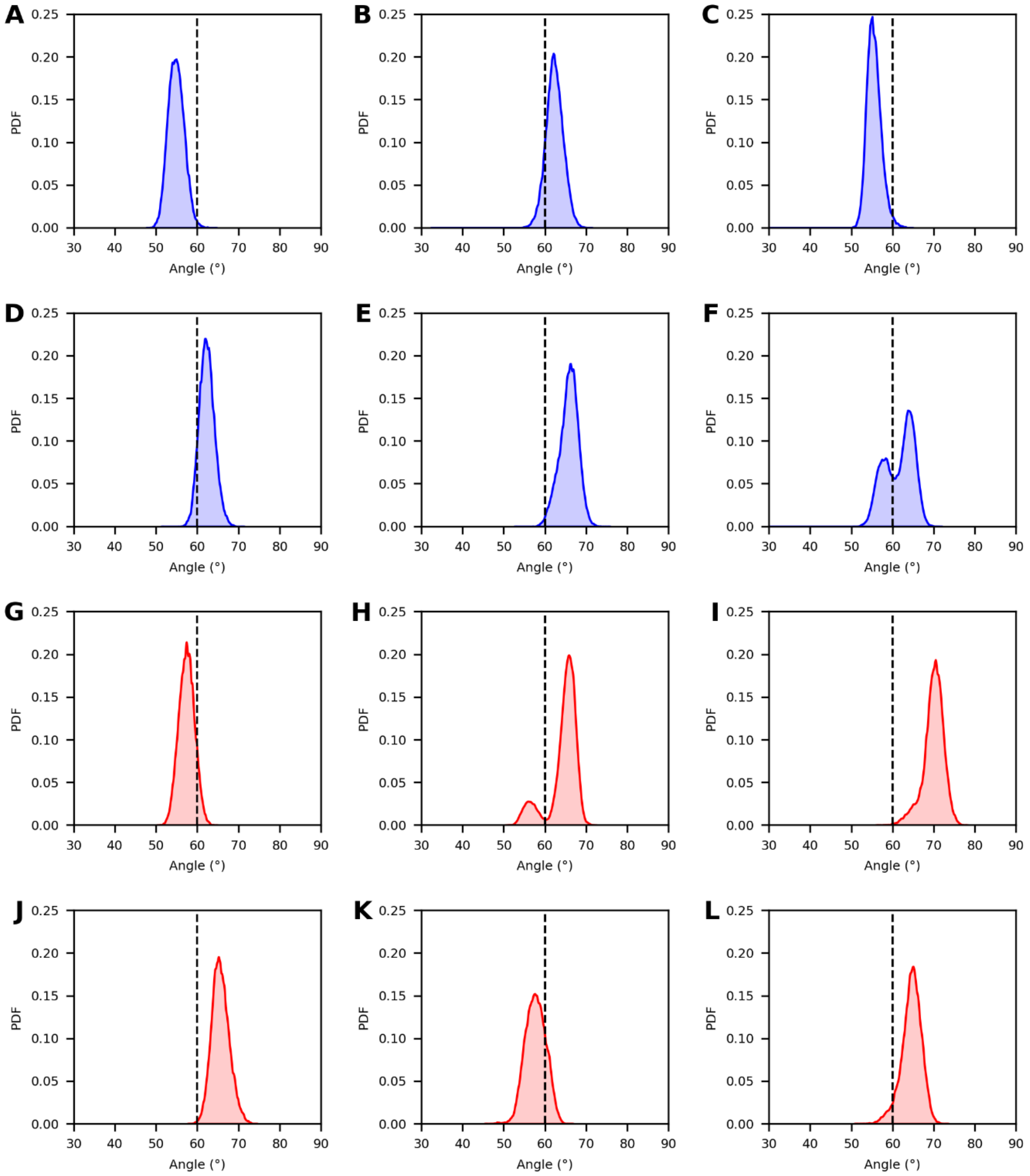


**Figure S20.** Distributions of NTH-TM1 angles in Cx32 wt:POPC:CHOL. **(A–F)** Angle distributions for each chain of the Cx32 hexamer during the first replicate MD simulation (blue). **(G–L)** Angle distributions for each chain during the second replicate MD simulation (red). Initial and final panels (A, F and G, L), and adjacent panels represent neighbouring chains in the structure. The vertical dashed line indicates the value of the angle in the cryo-EM structure of Cx32 in nanodiscs.

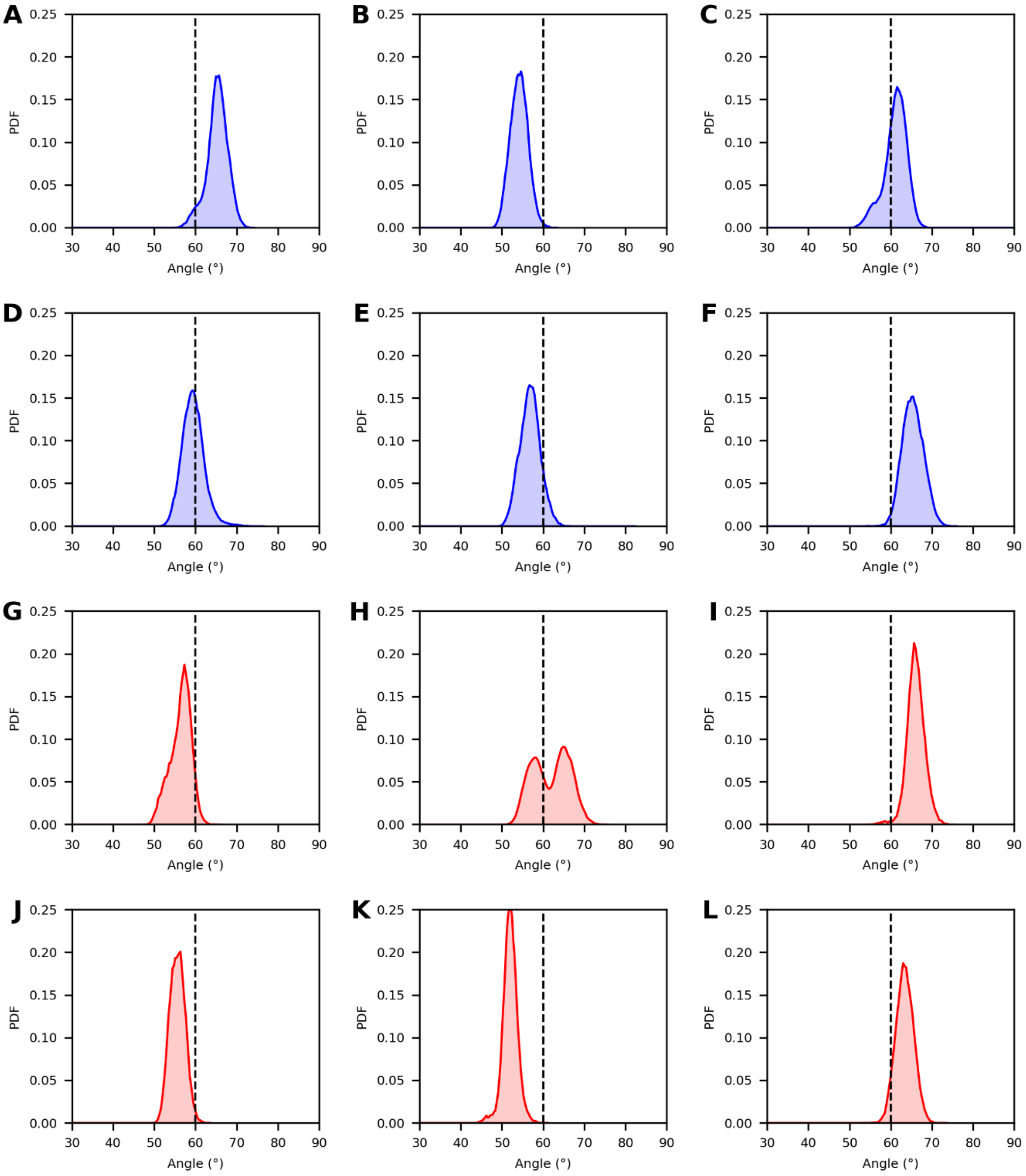


**Figure S21.** Distributions of NTH-TM1 angles in Cx32 W3S:POPC:CHOL. **(A–F)** Angle distributions for each chain of the Cx32 hexamer during the first replicate MD simulation (blue). **(G–L)** Angle distributions for each chain during the second replicate MD simulation (red). Initial and final panels (A, F and G, L), and adjacent panels represent neighbouring chains in the structure. The vertical dashed line indicates the value of the angle in the cryo-EM structure of Cx32 in nanodiscs.

**Table S1.** Cryo-EM data collection, data processing, and model building parameters.

| Data collection | | | | | |
|---|---|---|---|---|---|
| Sample | Cx32 GJC POPC | Cx32 GJC LPL | | W3S GJC POPC | Cx32 GJC no CHS POPC |
| Instrument | FEI Titan Krios/Gatan K3 Summit/Quantum GIF | | | | |
| Voltage [kV] | 300 | | | | |
| Electron dose [e-/Å] | 61.6 | Dataset 1: 61.2 | Dataset 2: 50 | 55 | 55 |
| Defocus range [µm] | -0.5 to -2.5 | | | | |
| Pixel size [Å] | 0.6506 | 0.6506 | | 0.6609 | 0.651 |
| Map resolution [Å] FSC 0.143 | 3.16 | 3.29 | | 2.35 | 3.12 |
| Number of particles | 80708 | 53450 | | 68649 | 15537 |
| Refinement | | | | | |
| Model resolution [Å] FSC 0.5 | 3.1 | 3.3 | | 2.3 | 3.1 |
| Map sharpening B-factor | -50 | -50 | | -20 | -20 |
| Map CC | 0.92 | 0.92 | | 0.91 | 0.89 |
| Model | | | | | |
| Protein residues/ligand | 2352/24 | 2352/24 | | 2316/0 | 2256/0 |
| ADP (B-factor), protein | 34.73 | 54.73 | | 68.25 | 85.96 |
| ADP (B-factor), ligand | 89.58 | 107.79 | | - | - |
| Bond length r.m.s.d. (Å) | 0.002 | 0.003 | | 0.003 | 0.003 |
| Bond angles r.m.s.d. (°) | 0.417 | 0.477 | | 0.480 | 0.474 |
| Validation | | | | | |
| MolProbity score | 1.12 | 1.52 | | 1.02 | 1.50 |
| Clash score | 2.86 | 4.34 | | 2.39 | 3.51 |
| Rotamer outliers (%) | 1.14 | 2.27 | | 0 | 2.96 |
| Ramachandran plot | | | | | |
| Favored (%) | 98.96 | 97.83 | | 98.63 | 98.37 |
| Allowed (%) | 1.04 | 2.17 | | 1.37 | 1.63 |
| Disallowed (%) | 0 | 0 | | 0 | 0 |
